# Impedance-based assay for pan-cancer early and rapid detection of cell-free DNA

**DOI:** 10.1101/2024.05.10.593096

**Authors:** Tejal Dube, Puja Prasad, Pragya Swami, Ankita Singh, Meenakshi Verma, Parul Tanwar, Shantanu Chowdhury, Shalini Gupta

**Affiliations:** Dept. of Chemical Engineering, Indian Institute of Technology Delhi, 110016, India; Amity Institute of Click Chemistry Research and Studies, Amity University, Noida, UP 201303, India; Integrative and Functional Biology Unit, CSIR Institute of Genomics and Integrative Biology (IGIB), 110025, New Delhi, India; Dept. of Pathology, Maulana Azad Medical College (MAMC), 110002, New Delhi, India; Academy of Scientific & Innovative Research (AcSIR), Ghaziabad, 201002, India; Asima Health Inc., 151 Charles St. West Suite #199, Kitchener, ON N2G 1H6, Canada

## Abstract

Aberrant DNA methylation is a hallmark of cancer, and plasma cell-free DNA (cfDNA) containing these abnormal methylation patterns has emerged as a promising non-invasive biomarker for pan-cancer detection. However, intrinsic challenges remain that continue to limit its broad clinical application. Here we show a simple and rapid impedance biosensor called Asima™ Rev that can detect cancer cfDNA in under 5 min without the need for any molecular labelling, electrode modification, signal amplification, or target enrichment steps. Using 216 clinical samples (50 healthy) from 15 different cancer types (all stages) we show an overall sensitivity and specificity of 96.4% and 94.0%, respectively. Differences in methylation content between cancerous and healthy cfDNA lead to distinct solvation behaviour and electro-physicochemical property that remain consistent across cancer types regardless of the distribution patterns of methyl cytosine. Our test exploits this inherent difference.

## INTRODUCTION

Liquid biopsy (LB) is a minimally invasive approach that has emerged as a promising tool for cancer screening and diagnosis through evaluation of blood-based biomarkers^1^. It currently commands a $4 billion global market share^2^. Circulating tumor cells (CTCs) and circulating tumor DNA (ctDNA) are the most widely studied LB biomarkers. ctDNA are tumor-derived fragmented genomic DNA (< 150 base pair long) that are released into the bloodstream from apoptotic or necrotic cancer cells. Their concentration levels thus increase in blood during cancer and can be used to track disease progression^3^. Normal population too contains a healthy baseline of cfDNA levels in their peripheral bloodstream (typically < 50 ng/ml), released as a consequence of the natural cell death in the body^4^. Concentration alone, however, is not a sufficient biomarker for cancer screening as the rate of ctDNA shedding depends on a number of different factors including the type, grade and stage of cancer^5^. Typical method for detecting ctDNA instead involves genomic mutational analysis using popular platforms like next-generation sequencing (NGS), digital or real-time PCR, mass spectrometry and microarray technology^6–9^. These platforms, however, have complex workflows, and require expensive infrastructure, limiting their clinical adoption^10^. Not only do they involve time-consuming library preparation, signal/target amplification steps and tedious data analysis, their clinical sensitivity also remains poor despite years of research and development^11^. This is mainly due to the low concentrations of ctDNA present in blood (< 0.01% in a pool of cfDNA background) and a limited number of recurrent somatic mutations present genome-wide, especially in early stages of the disease^12^. To date, there is no technique that can directly detect ctDNA from blood plasma at its originally low concentration, without the need for additional amplification steps.

More recently, companies like Grail and Singlera have expanded the horizon of cfDNA testing to include epigenetic signatures like methylation for multicancer screening^13^. Epigenetic remodelling in cancer involves both a global loss in methylation and tissue-specific repatterning across thousands of CpG dyad loci in the human genome^14,15^. While the coding and intergenic regions are extensively hypomethylated, the promoter regions of the CpG island clusters undergo a concomitant increase in methylation during cancer (**Fig. 1A**). This largescale cancer-specific redistribution of DNA methylation occurs early during carcinogenesis improving diagnostic accuracy compared to mutational analysis^16,17^. Moreover, this is a shared feature across most if not all cancer types, making cfDNA methyl landscape a universal biomarker for early cancer diagnosis^18^. The challenge with this approach, however, is that it relies on extensive clinical datasets and sophisticated machine learning tools for tissue of origin analysis, which contribute to its complexity and cost. The clinical performance is also sub-optimal. For instance, Grail’s Galleri test exhibits a low sensitivity of 51.5% and tissue of origin prediction accuracy of 88% despite conducting clinical trials in thousands of patients^19^. On the other hand, its specificity, indicating its ability to correctly identify non-cancerous cases, is notably high at 99.5%. It’s worth noting that sensitivity and specificity typically exhibit an inverse relationship (i.e., when one is low, the other is high), making it a significant technical challenge to achieve high values for both these parameters simultaneously.

**Fig. 1.**
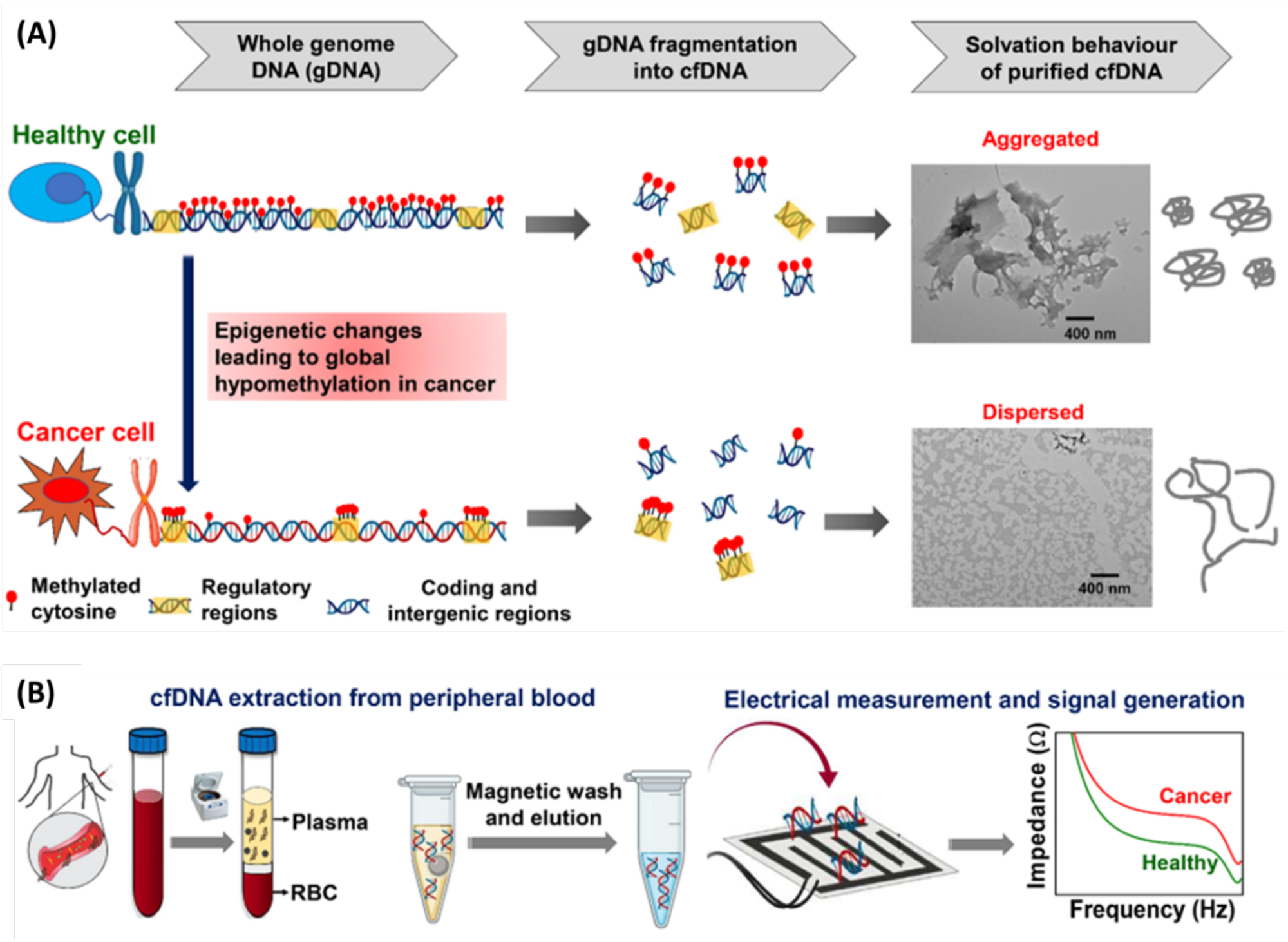
**(A)** Schematic depicting the epigenetic remodeling in genomic DNA that results into the differences in methylation patterns and solvation behavior of healthy and cancerous cfDNA. The TEM micrographs of purified cfDNA (70 ng/ml) taken from healthy (sample ID H11) and lung cancer (sample ID C98) samples showed healthy cfDNA to be more aggregated than cancer cfDNA. **(B)** Experimental workflow for Asima™ Rev for impedance-based cfDNA detection for pan-cancer screening.

Mainstream methodologies for DNA methylation analysis also involve sequencing (e.g., bisulfite, hydroxymethylcytosine, methylated DNA immunoprecipitation (MeDIP) *etc*.) which are long, costly, and cumbersome. Bisulfite conversion often leads to DNA fragmentation resulting in the generation of chimeric products^20,21^. Since circulating DNA is already short, further fragmentation during bisulfite treatment raises concerns, as it can potentially reduce the sensitivity of the assay. The variability in the quality of anti-5mC antibodies used in MeDIP can also affect the assay performance^22^.

In this paper, we introduce a new assay called Asima™ Rev that utilizes impedance spectroscopy to assess the global hypomethylation status of cfDNA. By measuring the electro-physicochemical changes influenced by epigenetic factors, our method offers a rapid and low-cost means for pan-cancer screening. In the past, Sina *et al*. have demonstrated the effect of epigenetic reprogramming on the physicochemical properties of DNA resulting in 3D conformational differences between healthy and cancerous DNA^23^. Inspired by this study, we hypothesized that genome-wide hypomethylation can also modify the solvation behavior of cancer cfDNA, allowing it to be distinguished from healthy cfDNA based solely on the difference in its conductivity in solution (**Fig. 1B**). To explore this hypothesis, we used a zwitterionic buffer that has previously been shown by our group to significantly enhance the impedance sensing capability of a biosensor^24^. This innovative approach enabled rapid differentiation of cancerous and healthy cfDNA within minutes, without the need of molecular labelling, electrode modification, sample preparation, signal amplification or target enrichment strategies. Unlike previously reported electrochemical sensors for cancer DNA (see **Table S1**), our impedance spectroscopy method represents a true one-step assay beyond the cfDNA extraction step.

In the following sections, we demonstrate the facile use of Asima™ Rev for multi-cancer screening in real world samples. We also investigate the role of several experimental parameters that effect the impedance response of both cancerous and healthy cfDNA samples. Additionally, we analyse the structural differences between cancerous and healthy cfDNA using transmission electron microscopy (TEM), revealing novel correlations with cancer stage. Finally, we look into the role of methylation in the molecular self-assembly of cfDNA molecules, by conducting experiments with differentially methylated oligonucleotides that mimic cfDNA molecules.

## RESULTS

To evaluate the clinical utility of our assay for pan-cancer screening, we tested a total of 216 clinical samples, comprising 166 cancer samples belonging to 15 different cancer types and spread across stages I to metastatic (see details in **Table S2**), and 50 healthy samples (see details in **Table S3**). Our experiments involved isolating the cfDNA fraction from human blood and suspending it in a zwitterionic buffer for impedance measurements. The electrical measurements were performed using off-the-shelf interdigitated electrodes connected to an impedance analyzer. No prior functionalization of the electrode substrate was required, and the cfDNA was measured directly at the concentration at which it was present in plasma, without any signal or target amplification. To minimize inter-assay variability in signals, every test measurement was preceded by the reading of a reference solution to carry out baseline correction. The final data were reported as the mod of differential impedance between the test and reference samples (|ΔZ| = |Z_cfDNA_ – Z_ref_|).

When plotted against cfDNA concentration, the impedance data at 100 kHz consistently exhibited smaller signals (lower |ΔZ| values) for cancer samples in comparison to healthy ones, regardless of the cfDNA concentration (**Figs. 2A,B)**. The binary classification threshold, determined using ROC analysis, was found to lie at -34 ohms, with an area under the curve (AUC) of 0.98 (**Fig. 2C**). The assay demonstrated an overall sensitivity (the fraction of true positives to all with disease) of 96.39% and specificity (the fraction of true negatives to all without disease) of 94.00% across all cancer types. The sensitivity values improved with the stage of cancer, with stage I samples showing 88% sensitivity versus metastatic samples showing 100% (**Fig. 2D**). No discernible patterns in signals were observed as a function of cancer type (**Fig. 2E**). The concentration of cfDNA alone was not found to be a reliable classifier, in agreement with the literature (**Fig. 2F**)^25,26^. The TEM analysis of kidney cancer (**Figs. 2G-top, S1**), head and neck cancer (**Figs. 2G-bottom**, S1), and breast cancer (**Fig. S2**) samples revealed fascinating trends of reduced cfDNA aggregation with advancing cancer (stages I to IV), despite originating from different patients.

**Fig. 2.**
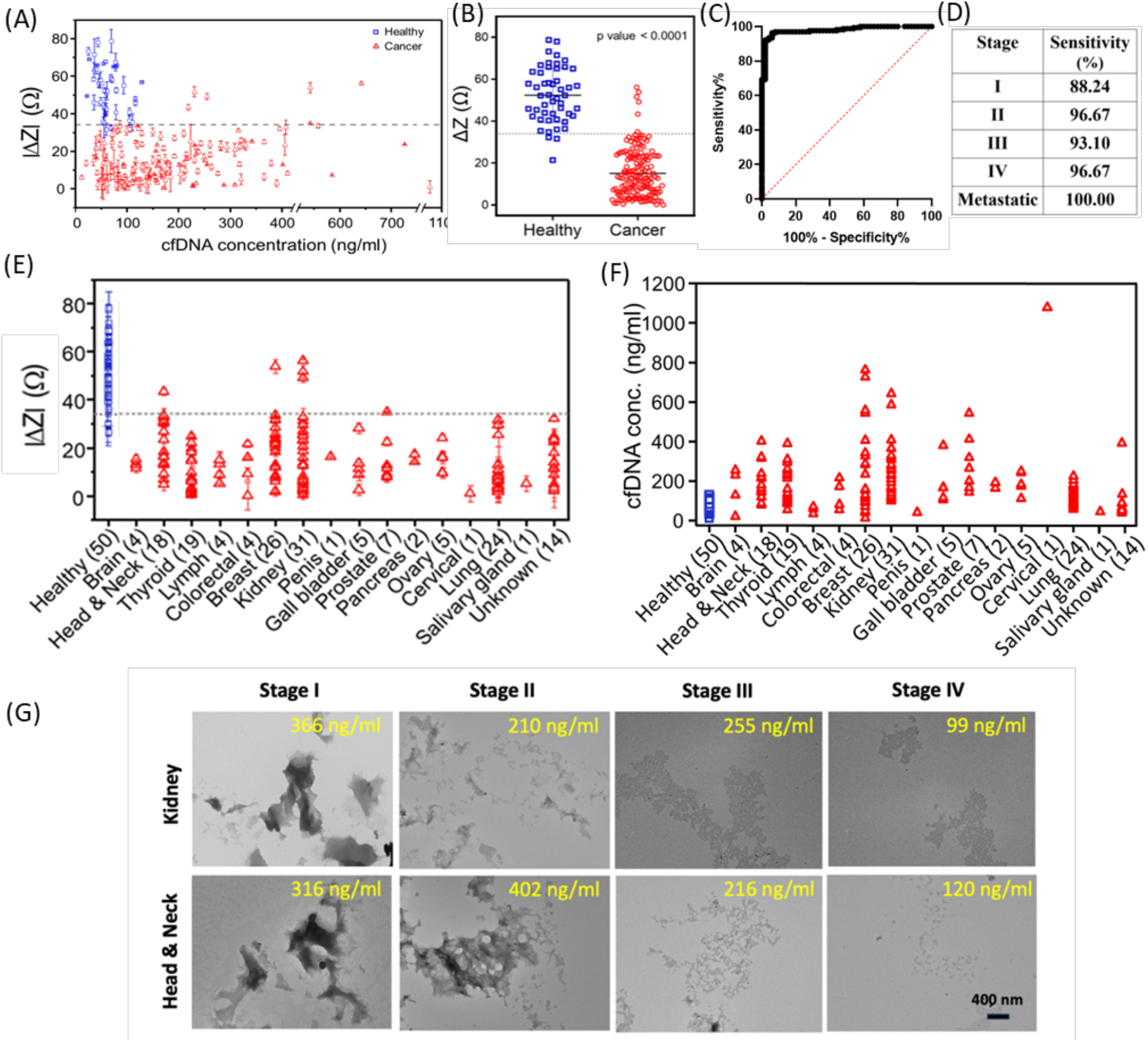
Clinical assay validation performed on 50 healthy and 166 cancer samples. **(A)** Differential impedance response as a function of cfDNA concentration showing cancer samples falling below the threshold (indicated using a dotted line at -34 ohms), while healthy samples lay above it. **(B)** Unpaired Student’s t-test indicating high statistical significance (p-value < 0.0001) between healthy and cancer data at the 95% confidence interval. **(C)** ROC analysis with AUC of 0.98. **(D)** Sensitivity data for cancer staging at the population level. **(E & F)** Correlation of impedance change (E) and cfDNA concentration (F) in 15 different types of cancers. The dotted line in (E) indicates the -34 ohms threshold. (**G**) TEM images of kidney (L-R: C62, C65, C74, C79), and head and neck (L-R: C41, C45, C48, C53) cancer samples at different cancer stages.

To understand the origins of these signal differences between healthy and cancerous cfDNA, we carried out a detailed study to investigate the role of experimental factors like the applied electric field, liquid medium in which the cfDNA is suspended, and the concentration and structure of cfDNA, that can influence the impedance response of the system. Our findings are discussed below.

### (i) Effect of electric field

The frequency-dependent impedance spectra of healthy and cancer cfDNA samples showed a plateau (**Fig. 3A**) and low phase angle (θ < -2°) (**Fig. 3B**) between 10 kHz to 1 MHz, suggesting that the system behaves mainly as a resistor in this frequency range^27^. The healthy cfDNA was found to be more conductive than cancer cfDNA (lower Z value) at all frequencies. The difference in signal between healthy and cancer samples was found to be highest at 100 kHz (ΔZ most negative), so this frequency was prioritized for data collection. The effect of voltage was found to be invariant in the 5 to 500 mV range suggesting negligible Faradaic or Joule effects (**Fig. S3**).

**Fig. 3.**
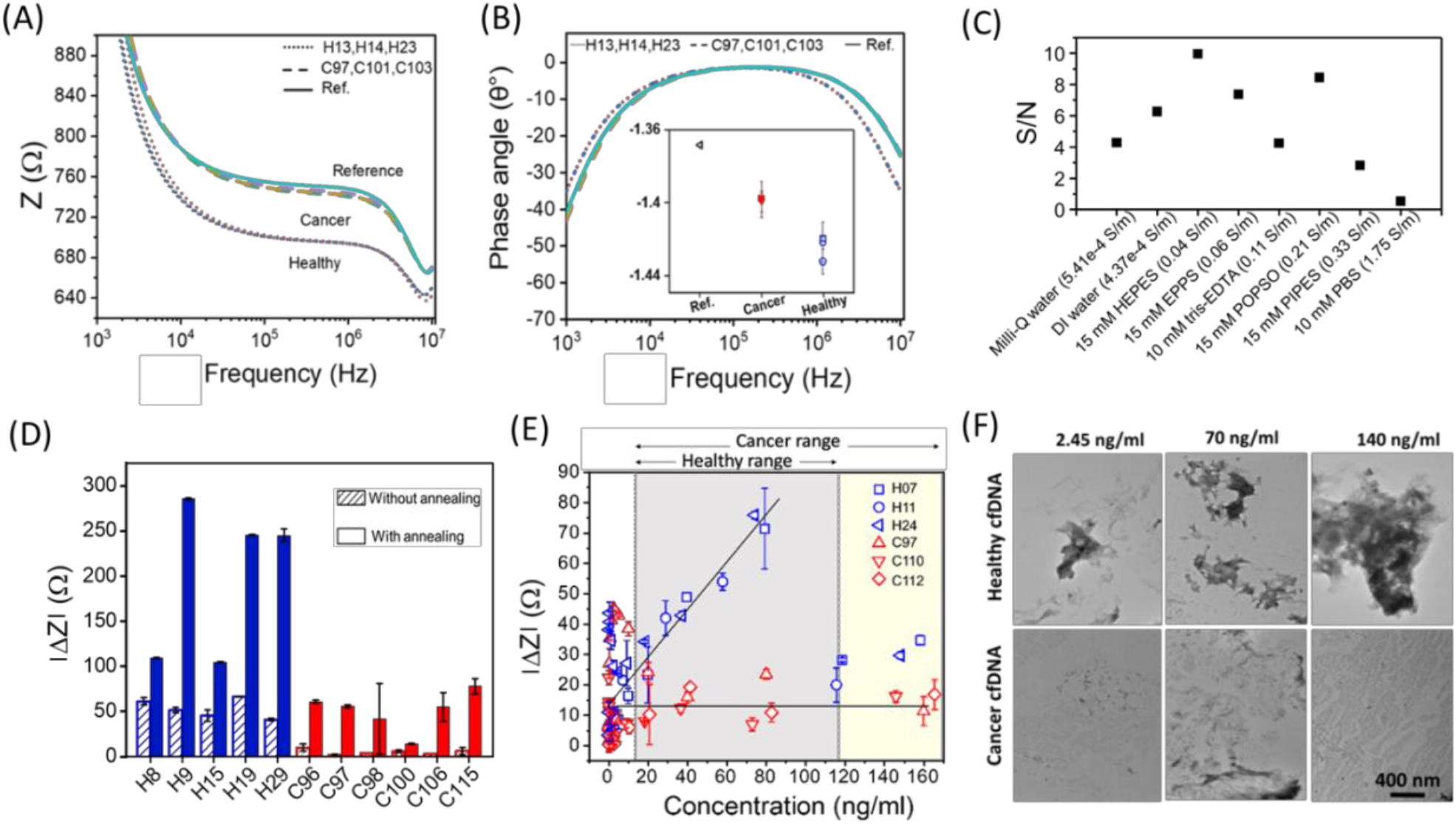
Frequency-dependent impedance spectra **(A)** and phase angle **(B)** of reference solution, healthy cfDNA samples (H13, H14, H23) and cancerous cfDNA samples (C97, C101, C103). The inset in (B) shows phase angles obtained at 100 kHz. **(C)** Effect of liquid medium on impedance response showed HEPES buffer to have best performance (highest S/N). The data are arranged in increasing order of medium conductivities and measured in sample ids H25 and C104 (see SI for sample calculations). **(D)** The effect of cfDNA annealing on impedance response in healthy (H8, H9, H15, H19, H29) and cancerous (C96, C97, C98, C100, C106, C115) cfDNA samples. (**E**) The effect of cfDNA concentration studied through serial dilution of healthy (H07, H11, H24) and cancer (C97, C110, C112) cfDNA samples. The results were highly non-monotonic (see Fig. S4) for zoomed-in plots at lower concentrations). The lines are drawn to aid the eye. **(F)** TEM images showing higher aggregation with increasing concentration in both healthy (H11) and cancer (C98) cfDNA samples. Error bars where not visible are smaller than the data points. All data are obtained at 100 kHz where not explicitly mentioned.

### (ii) Effect of medium

Next, we studied the effect of medium. As the electrical conductivity of cfDNA is measured in solution, the type of medium present in the background can significantly influence the signal quality. Based on our past research, zwitterionic buffers show significantly superior performance in enhancing impedance signal sensitivity compared to simple electrotypes, owing to their (i) low intrinsic conductivity, (ii) low concentrations used in biological experiments, (iii) low ionic dissociation constants and (iv) high ionic-ionic interactions^24^. In the present case too, HEPES showed the best performance (highest signal-to-noise ratio) compared to all the other media studied, whereas phosphate buffered saline (PBS) was found to be the worst (**Fig. 3C & Table S4**).

### (iii) Effect of annealing

Annealing is a crucial parameter, known to significantly impact the electrical conductivity of DNA samples. This process is widely acknowledged for its role in enabling DNA to adopt a more structured and stable conformation, leading to enhanced stacking interactions between base pairs and increased π-π electron overlap. Consequently, annealing leads to an overall increase in DNA conductivity^28^. In our experiments too, we observed a significant increase in cfDNA conductivity following the annealing step in all the cases (**Fig. 3D**) (note here that ΔZ value is actually negative for every data point implying that samples are more conductive than reference solution). More importantly, the relative difference in signal between healthy and cancer samples improved by more than 3-fold post-annealing (**Table S5**), suggesting that this could be used as an additional parameter for enhancing the assay sensitivity.

### (iv) Effect of cfDNA concentration

We further investigated the change in impedance signal with respect to cfDNA concentration. To evaluate this parameter, a higher than usual stock concentration of healthy and cancer cfDNA samples was prepared and sequentially diluted to lower concentrations reaching into the sub-clinical range. The clinical concentration ranges were identified based on the values reported in literature, however it’s worth noting that the literature itself presents a considerable variation in these ranges.^4,29^ The impedance values obtained from these samples exhibited a highly non-monotonic behaviour with respect to concentration (**Fig. 3E, S4**). Specifically, the curve for cancer cfDNA showed a single peak in the sub-clinical concentration range, whereas the data for healthy cfDNA showed two peaks, one within the sub-clinical range and another notably higher peak towards the upper end of the clinical concentration range. Within the clinical range, cancer cfDNA samples consistently showed lower |ΔZ| values compared to healthy samples. The lowest limit of detection (LoD) was found to be 0.4 ng/ml.

TEM visualization of cancerous and healthy cfDNA samples across various concentrations revealed that healthy cfDNA samples exhibited considerably higher aggregation across all the concentrations, whereas cancerous cfDNA samples remained dispersed throughout (**Figs. 3F, S5, S6**). The extent of aggregation appeared to increase with concentration, particularly evident in healthy samples, where micron-sized amorphous DNA networks were observed at the highest concentration of 140 ng/ml. Drawing correlations between these TEM findings and our impedance data, we inferred that the differential impedance values (|ΔZ|) increase with aggregation in the clinical range and can be utilized as a valuable means of distinguishing between cancerous and healthy samples based solely on their morphological status and, consequently impedance measurements.

### Impedance response of differentially methylated oligonucleotides

Finally, to confirm the role of methylation in the architectural and impedance variation between healthy and cancer cfDNA samples, we conducted experiments with synthetic oligonucleotides that were specifically designed to mimic the cancer-induced methylation patterns in cfDNA. The 77-mer long oligonucleotide sequences, chosen from the hTERT gene promoter region, each had a different extent of methylation content as shown in **Table 1**^30^. Both single-stranded (ssDNA) and double-stranded (dsDNA) oligonucleotides were studied as the cfDNA is known to exist in both forms^31^. The impedance results showed that |ΔZ| increased monotonically as the methylation content was increased from 10% to 100% at a fixed molecular concentration in both ssDNA and dsDNA (**Fig. 4A**). This corroborated with our cfDNA results, which also displayed higher |ΔZ| values in healthy samples having higher hypermethylation content than cancer cfDNA. The phase angle data were also found to be consistent with Fig. 3B results, with greater shifts observed in more methylated samples (**Fig. 4B**). The dsDNA samples were consistently found to be more conductive than ssDNA samples corroborating with the existing literature^32^.

**Table 1.**
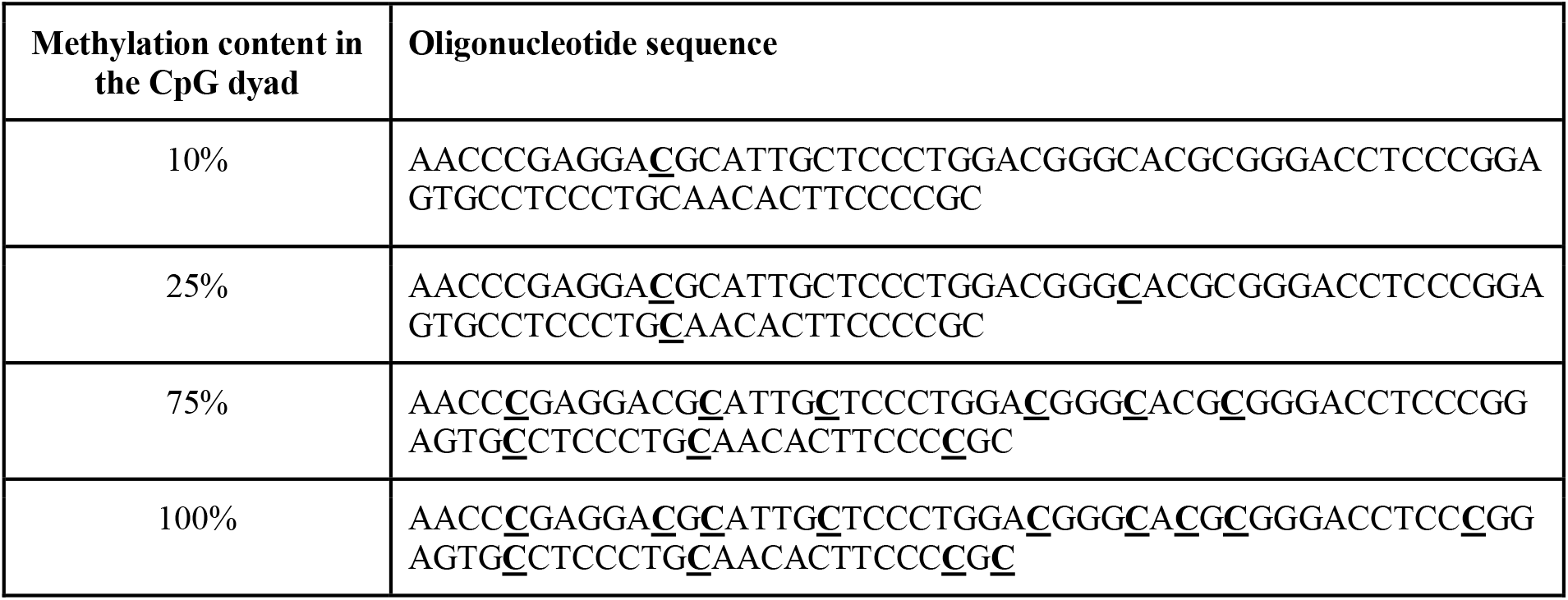
The oligonucleotide sequences used in our experiments.

**Fig. 4.**
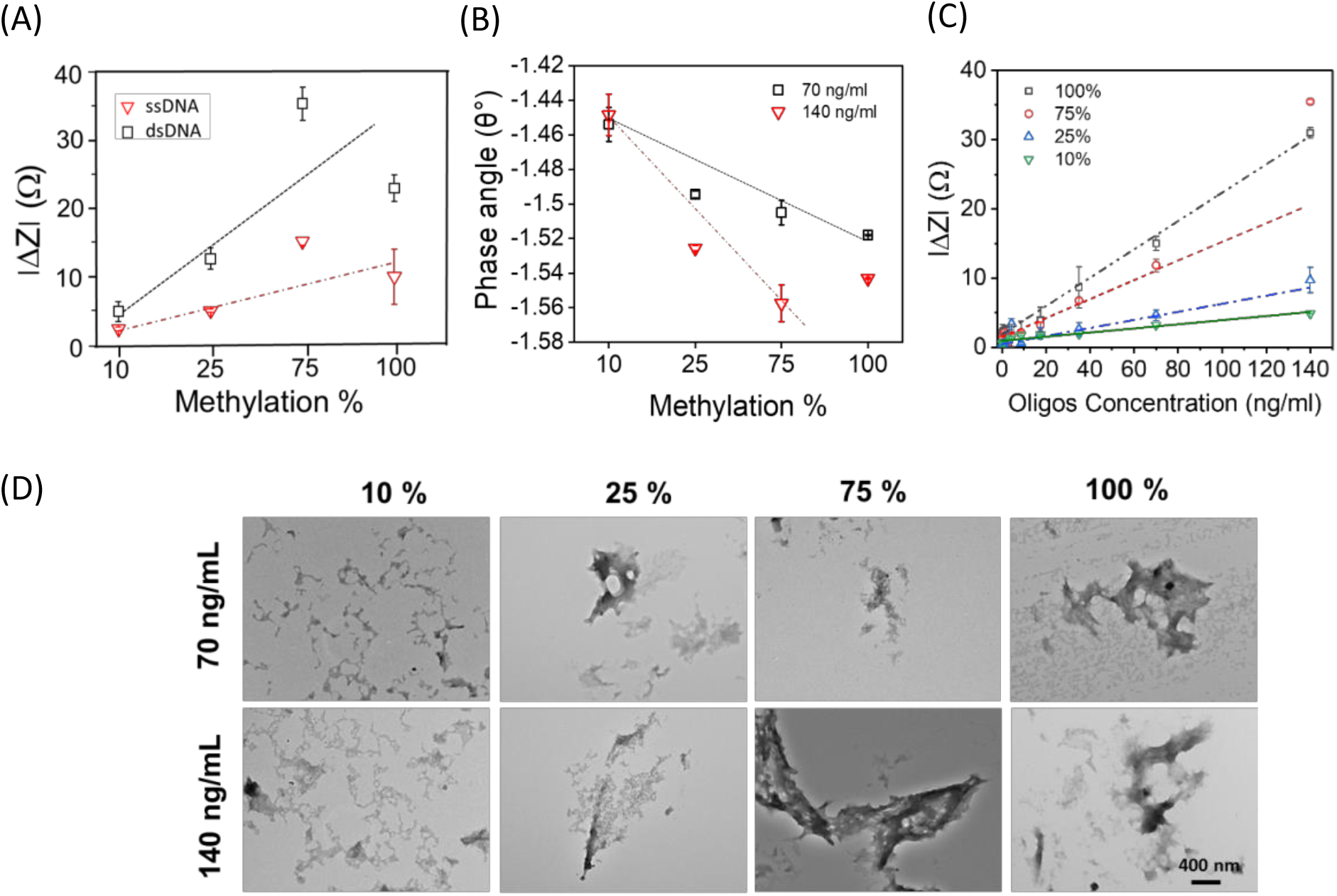
Results with differentially methylated oligonucleotides. **(A)** Change in ssDNA and dsDNA impedance response with increasing methylation content at a fixed oligo concentration (140 ng/ml). **(B)** The change in phase angle of dsDNA as a function of methylation content at two different concentrations. **(C)** The change in impedance response of differentially methylated dsDNA taken at different concentrations. The lines are drawn to aid the eye. Error bars where not visible are smaller than the data points. **(D)** Representative TEM micrographs of 10% to 100% methylated dsDNA at two different concentrations (70 and 140 ng/ml) illustrating the propensity of molecules to aggregate when there is higher methylation content in the system. The scale bar is 400 nm in all the images.

Further experimentation involved varying the methylation content through oligonucleotide concentration instead of the extent of methylation. The data were collected for various differentially methylated oligos and the results obtained illustrated expected trends in both dsDNA (**Fig. 4C)** and ssDNA (**Fig. S7**), with the rate of signal change varying proportionally to the methylation content. Our TEM data also supported our impedance findings, with dsDNA samples containing higher methylation content (through both higher extent of conjugation and oligo concentration) exhibiting greater aggregation (**Figs. 4D, S8**). Collectively, these results provided compelling evidence that difference in methylation content is the predominant factor driving signal change in our assay, confirming the critical role of methylation in distinguishing between healthy and cancerous cfDNA.

## DISCUSSION

Aberrant DNA methylation is a hallmark of oncogenesis, and differences in methylation patterns between tumor and benign tissues have been reported in many cancer types^18^. A characteristic feature of epigenetic remodelling in cancer gDNA is the differential hypomethylation in the coding and intergenic regions and differential hypermethylation in the CpG-rich regulatory regions such that the overall genomic landscape is markedly hypomethylated^17,23^. A recent study also found that cfDNA fragments derived from gDNA are selectively enriched in coding and intergenic regions compared to the gene promoter regions in both malignant and benign samples^33^. These findings imply that the structural variations in cancer and healthy cfDNA seen in our experiments can arise from the differences in their methylation patterns, with healthy cfDNA containing significantly higher number of methyl groups as depicted in Fig. 1A. It is the presence of these non-polar functional groups that likely leads to the hydrophobic self-assembly of healthy cfDNA in aqueous solution. As the loss of methyl groups from the cytosines happens gradually during cancer^34^, the morphology changes from microaggregates to dispersed nanoaggregates as the cancer progresses, which is made evident by the stage-wise TEM images of the cfDNA samples shown in Fig. 2G. More importantly, since the global methylation loss is a common signature shared by multiple cancer types^35^, the electro-physicochemical properties are found to be similar across various cancer types.

Methyl-induced aggregation can also profoundly impact impedance data in an alternating current electric field. The cfDNA concentration-dependent non-monotonic behavior observed in our experiments can be explained as illustrated in **Fig. 5**. Here, in the very low concentration regime (section a), cfDNA molecules are relatively well-dispersed and isolated from each other, such that the positively charged counterions along their phosphate-rich backbone can freely move under the perturbation of an external electric field. As the number of base pairs per unit volume is increased, this results in a greater charge density (where mobile charges ∝base pairs) and higher |ΔZ| values, i.e., solution becomes more conductive. When the cfDNA concentration is increased a little further approaching section b, the |ΔZ| values start to drop due to the electrokinetic overlap between isolated molecules or very small assemblies. This type of decrease in conductivity has been previously reported by our research group for gold and polystyrene colloidal particles^36^ and is unique to zwitterionic buffers due to their special properties^24^. Also, the peak was found to shift to lower packing fractions for smaller particles. In the present case of cfDNA, the “particle” is ∼ 1 nm (few molecules), which explains the first peak at several orders (∼10^5^) lower volume fraction as compared to our previous study. The slight discrepancies observed between the actual experimental data of healthy and cancerous cDNA in this low concentration regime (refer to Fig. S4) may arise from the differences in their length, given that cancer-derived cfDNA is shorter than healthy-derived cfDNA^37^ and conductivity is directly proportional to DNA length^38^.

**Fig. 5.**
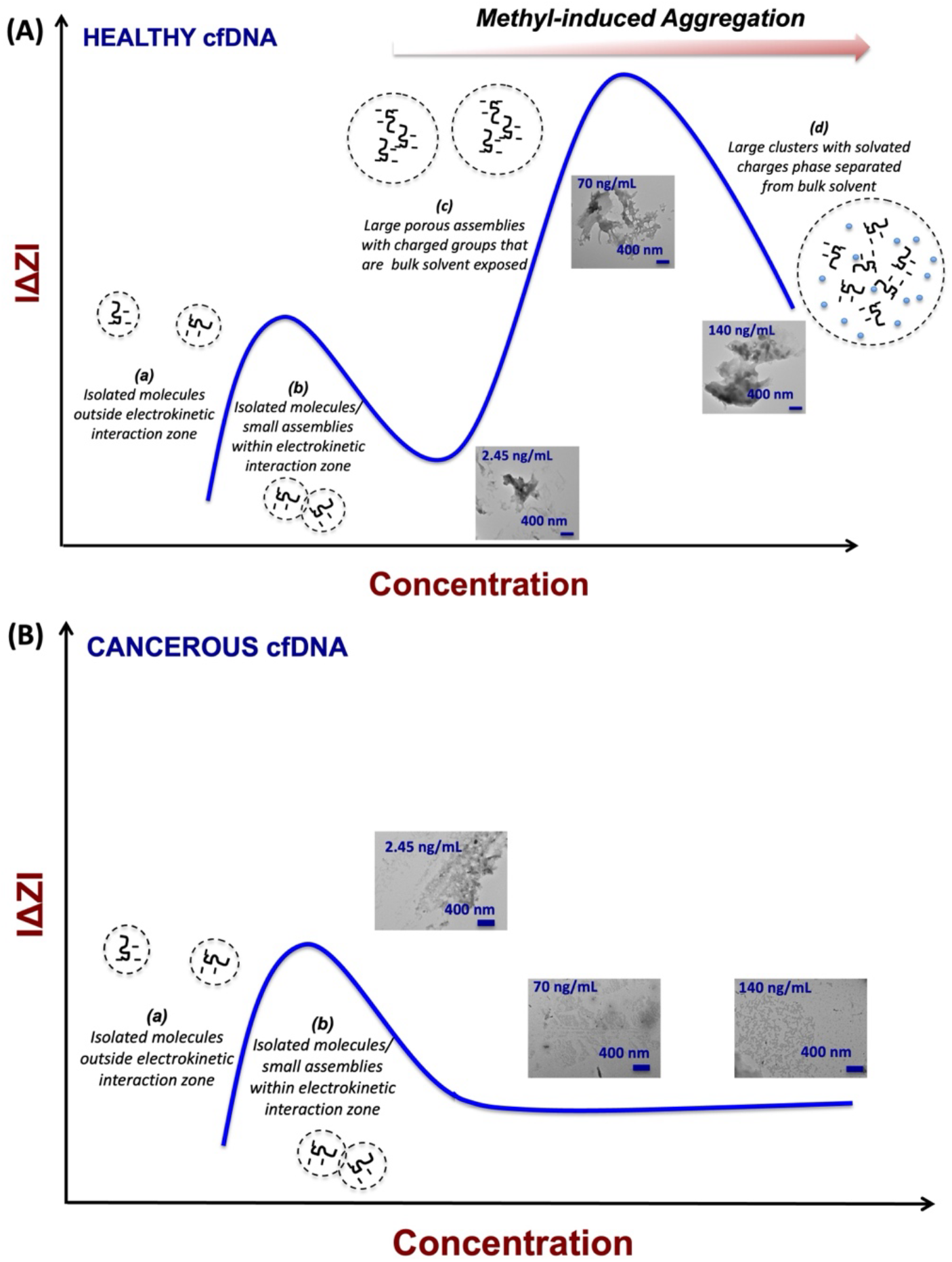
The proposed mechanism for differential impedance variation in healthy **(A)** and cancerous **(B)** cfDNA samples as a function of concentration. In healthy cfDNA, the initial peak reflects the electrokinetic overlap between individual molecules or tiny assemblies. This overlap radius is indicated by the dotted line surrounding the cfDNA molecules. Following this peak, there is an increase in conductivity, attributed to the formation of porous assemblies driven by hydrophobic interactions among methyl groups. These assemblies reduce the electrokinetic overlap while maintaining contact with the surrounding solvent through charged groups. The second peak signifies the emergence of larger clusters where the charged groups are removed from the bulk solvent, leading to a decrease in conductivity. Conversely, in cancerous cfDNA, individual molecules are consistently present in sufficient numbers, barring extremely low concentrations, thus the electrokinetic overlap remains largely unaltered, resulting in a plateau beyond the initial peak.

At intermediate concentrations of healthy cfDNA (section c in Fig. 5A), large porous clusters start to emerge reducing the “particle” concentration and thus, electrokinetic overlap. This is accompanied by an increase in |ΔZ| values (solution becomes more conductive) as the clustering leads to the exposure of charged groups to the bulk solvent. Such conductivity enhancement due to charge percolation effects has been previously reported for other nanoscale colloidal systems^39^. In this regime, annealing can improve the packing to increase solvation of charged groups by bulk water and ions (thermodynamically favourable state), leading to significant conductivity increase as seen in Fig. 3D. Since the aggregation is largely methylation-induced, the second peak is only seen in case of healthy cfDNA whereas, for cancer cfDNA, there is only one dominant mechanism, involving overlap of electrokinetic interaction zones of approximately isolated molecules or small assemblies. Therefore, this data shows only one peak and the signal remains largely invariant after the formation of nanoassemblies, despite the increase in cfDNA concentration (Fig. 5B). At very high concentrations (section d in Fig. 5A), poorly soluble dense particle clusters of healthy cfDNA are formed and removed from the bulk due to settling and/or phase separation of solvated charges from the bulk solvent^39^. This again leads to a decline in conductivity.

It is noteworthy that maximum signal variation between healthy and cancer samples occurs within the clinically-relevant concentration range which is a crucial aspect for our assay, as it offers a rapid way of identifying cancer samples. The parallel trends observed between cfDNA molecules and synthetic oligos imply that our measurements are influenced by epigenetic modifications. The higher conductivity in dsDNA compared to ssDNA is attributed to its inherently more stable structure that fosters increased stacking interaction between the base pairs and greater π-π electron overlap.

## CONCLUSIONS

In summary, we demonstrate a simple and rapid impedance biosensor for cfDNA methylation analysis for pan-cancer screening. Using 216 samples, we show relatively high sensitivity and specificity of detection for Asima™ Rev compared to other systems. Mechanistically, this is linked to solvation differences induced by altered methylation state of normal versus cancer cfDNA isolated from blood. In the past, electrical biosensors have been widely used for DNA characterization due to their robustness, ease of miniaturization, low detection limits, and compatibility with biological fluids^40^. Traditionally, DNA detection by integrated sensors requires the molecule to be immobilized on the electrode surface using a bioaffinity receptor followed by its signal amplification using a set of probes or chemically active compounds. This not only adds extra steps and cost of reagents but also increases the chances of false positives due to non-specific adsorption of the molecules on the surface. The electrode design/material in turn needs to undergo extensive optimization for the various binding steps and the chips cannot generally be reused, making the process less eco-friendly and economically inefficient. Some recent studies have tried to address these issues by measuring the electrical properties of gDNA directly in solution, however, their signal sensitivity is low and greatly influenced by the type of solvent used. In deionized water for example, the lower LoD was found to be > 10 mg/ml for gDNA^41^. However, no such example exists for cfDNA. Our biosensor is the first of its kind to simplify both the test preparation and the sample measurement steps. The test is PCR-free, does not require sequencing infrastructure and gives ready readout. Together, these present a novel and simple biomarker analysis that is rapid, can be readily accessible, and equitable.

## MATERIALS AND METHODS

### Materials

Platinum-on-glass planar interdigitated electrodes (IDEs) with 40 symmetrical fingers (20 pairs) of 3.5 mm length and 100 μm width each, and 100 μm edge-to-edge spacing (Micrux Technologies, Spain); Sodium hydroxide (NaOH) (Merck, India); Acetone (Fisher Scientific); MagMax cell-free DNA isolation kit (A29319, Thermofisher); Phosphate buffer saline (PBS), 4-(2-hydroxyethyl)-1-piperazineethanesulfonic acid (HEPES), 4-(2-hydroxyethyl)-1-piperazinepropanesulfonic acid (EPPS), piperazine-1,4-bis(2-hydroxypropanesulfonic acid) (POPSO), piperazine-N,N′-bis(2-ethanesulfonic acid) (PIPES), tris-HCl buffer, ethylenediamine tetra-acetic acid (EDTA), and custom-synthesized differentially methylated oligonucleotides shown in Table 1 (Sigma-Aldrich); Polydimethylsiloxane (PDMS) elastomer kit (Dow Corning, USA); K3EDTA blood collection tubes (Greiner Bio One India Pvt. Ltd.); 200 mesh square carbon TEM grids (catalogue CF200-CU, Electron Microscopy Sciences); Cryovials (Tarson). All buffers were prepared at pH 7.4 in ultrapure deionized Milli-Q water (∼18.2 MΩ.cm resistivity) following standard procedures and experiments were conducted at room temperature (RT ∼ 25 °C).

### Methods. (i) Sample collection

A total of 166 cancer samples from newly diagnosed patients with no prior treatment were collected from Maulana Azad Medical College and Rajiv Gandhi Cancer Institute and Research Centre in Delhi after obtaining ethics approval (F.1/IEC/MAMC/67/02/2019/No.141 and Res/BR/TRB-19/2020/56). These cancer samples belonged to 15 different cancer types spread across different stages in males and females between 20-90 yrs of age group (mean age 54.42 yrs; cfDNA concentration 12-1077.41 ng/mL; see Table S2 & **Fig. S9** for details). The healthy samples were collected from a mix of 50 male and female volunteers in a similar age group after obtaining informed consent (mean age 34.76 yrs; cfDNA concentration 22-129 ng/mL; see Table S3 & Fig. S9 for details). From each participant, 4 ml whole blood was collected in a K3EDTA tube via routine venous phlebotomy and the plasma was isolated from blood by centrifugation at 2000g for 10 min at 4°C, within 4 h of blood collection. The resulting plasma was again centrifuged at 16000g for 10 min at 4°C, and the supernatant was collected and transferred into cryovials before immediately freezing at -80°C.

### (ii) cfDNA extraction

Plasma samples at -80 °C were sequentially thawed, first at -20 °C for at least 2 h and then at RT for 1 to 2 h. The cfDNA was extracted using the MagMax™ cfDNA isolation kit following a slightly modified protocol from the manufacturer. Briefly, plasma was treated with magnetic beads and the lysis/binding buffer. The cfDNA-bound magnetic beads were then separated using a magnetic stand followed by washing with the wash solution and final washing with 80% ethanol. The cfDNA was finally eluted in 20 μl of elution buffer (henceforth referred to as the “stock”) and used for preparing all the test samples after appropriate dilution. The cfDNA concentration in the stock was quantified using Nanodrop™ (Lite, Thermo Fisher Scientific).

### (iii) cfDNA sample preparation

A required volume of cfDNA solution was taken from the stock and topped with the elution buffer to make the total volume 2 μl. This 2 μl volume was then spiked into 98 μl of 15 mM HEPES buffer at pH 7.4 (or any other medium depending on the study) to get the final test sample volume. Unless otherwise stated, the stock volume was always chosen such that the cfDNA concentration in the final test sample was equal to that in plasma (see sample calculation in SI). For cfDNA annealing experiments, the cfDNA samples were prepared as mentioned above and their impedance response was recorded as discussed later. Following this, a 40 μl aliquot of the same sample was taken, along with a 40 μl aliquot of reference solution (2% v/v elution buffer in 15 mM HEPES buffer pH 7.4), and the tubes were kept in a dry heat bath at 95 °C. After 5 min of heating, the samples were removed and allowed to cool at RT to compare the effect of annealing. To study the effect of cfDNA concentration, we started the cfDNA extraction process using a larger plasma volume and eluted the cfDNA in 20 μl elution buffer. This highly concentrated stock was then serially diluted with the reference solution to the get the final sample concentrations.

### (iv) Oligonucleotide sample preparation

ssDNA powder was reconstituted in 10 mM tris-HCl buffer at containing 1 mM EDTA to prepare a 100 μM stock. The stock concentration was converted to ng/μl using Nanodrop™ and the 100 μM stock was diluted to a working stock of 7 μg/ml using 10 mM tris-HCl buffer containing 1 mM EDTA. The final test sample of 140 ng/ml was prepared by mixing 2 μl of working stock with 98 μl of 15 mM HEPES buffer. Similarly, the final test sample of 70 ng/ml concentration was prepared by mixing 1 μl of working stock with 1 μl of tris-HCl buffer containing 1 mM EDTA and 98 μl of 15 mM HEPES buffer. For dsDNA, equimolar concentrations (100 μM each) and equal volumes (2.5 μl each) of complementary ssDNA were taken from the stock and hybridized by heating the mixture at 95 °C for 5 min followed by slow cooling at RT. The concentration of this hybridized solution was noted using Nanodrop™ and used to prepare a working stock of 7 μg/ml in 10 mM tris-HCl buffer containing 1 mM EDTA. The final test sample with 140 ng/ml concentration was prepared by mixing 2 μl of working stock with 98 μl of 15 mM HEPES buffer. The 70 ng/ml concentration test sample was prepared by mixing 1 μl of working stock with 1 μl of tris-HCl buffer containing 1mM EDTA and 98 μl of 15 mM HEPES buffer.

### (v) TEM characterization

cfDNA and oligonucleotides were characterized using TEM (JEOL JEM-1400). For this, 10 μl of each sample was spotted dry onto a TEM grid, and stained with 1% uranyl acetate. Images were then taken at 120 kV at different magnifications.

### (vi) PDMS microchamber preparation

A solidified PDMS sheet of 1.6 mm thickness was prepared using a standard protocol^42^. The PDMS sheet was cut into multiple pieces of 1.5 cm x 1.2 cm size each. A 4 mm diameter hole was punched through each PDMS piece using a standard paper puncture to create a microchamber. The pieces were then covered by scotch tape on both sides and placed in a clean dry box at RT until further use. At the end of each experiment, the PDMS microchamber could be reused after sequential washing with 15 mM HEPES buffer, 1 M NaOH, MilliQ-water, and acetone. The chamber was dried and stored at RT after covering it with scotch tape on both sides.

### (vii) Impedance measurements

Before each experiment, the scotch tape was removed from both sides of the PDMS and the microchamber was placed on top of a clean pair of IDEs. Twenty microliters of reference solution was then added into the PDMS chamber and the top was covered with a glass coverslip to avoid evaporation (see **Fig. S10**). The sample was allowed to stabilize for 3 min and its impedance reading was then recorded at 100 mV and 20 Hz to 2 MHz frequency range using an impedance analyser (Keysight E4990A). The process was repeated with the test sample and the final result was reported as the mod of differential impedance (|ΔZ| = |Z_test_ – Z_ref_|) at 100 kHz. All experiments were performed in duplicate using independently prepared test samples from the same stock and the data were plotted as the mean of signal ± 2 SD. Where possible, unpaired Student’s t-tests was carried out between independent data sets using GraphPad Prism 8 software to test their statistical significance using the null hypothesis (*p*). The confidence interval for rejecting the null hypothesis was chosen as 95% (*p* ≤ 0.05). GraphPad Prism was also used to calculate the receiver operating characteristic (ROC) curve for sensitivity and specificity analysis.

At the end of each experiment, the IDEs were washed with 15 mM HEPES at least thrice or until the expected impedance reading was obtained with HEPES. The electrodes were then dried and stored in a clean dry box at RT until further use. If the impedance reading was not restored despite several washes (happened approximately after 50 experiments or so), the electrodes were restored after washing with 1 M NaOH and this electrode could be indefinitely used if maintained well (**Fig. S11**).

## Acknowledgements

SG is grateful to Prof. Gaurav Goel for insightful discussions on the role of cfDNA concentration in impedance data analysis. PP acknowledges CSIR for SRA position (Pool No. 9031-A) and SERB-DST (SRG/2023/001099) for SRG grant.

## References

1. Marrugo-Ramírez, J., Mir, M. & Samitier, J. Blood-Based Cancer Biomarkers in Liquid Biopsy: A Promising Non-Invasive Alternative to Tissue Biopsy. Int. J. Mol. Sci. 19, 2877 (2018).

2. Bargahi, N., Ghasemali, S., Jahandar-Lashaki, S. & Nazari, A. Recent advances for cancer detection and treatment by microfluidic technology, review and update. Biol. Proced. Online 24, 5 (2022).

3. Das, J. et al. An electrochemical clamp assay for direct, rapid analysis of circulating nucleic acids in serum. Nat. Chem. 7, 569–575 (2015).

4. Alborelli, I. et al. Cell-free DNA analysis in healthy individuals by next-generation sequencing: a proof of concept and technical validation study. Cell Death Dis. 10, 534 (2019).

5. Siravegna, G. et al. How liquid biopsies can change clinical practice in oncology. Ann. Oncol. 30, 1580–1590 (2019).

6. Heitzer, E., Haque, I. S., Roberts, C. E. S. & Speicher, M. R. Current and future perspectives of liquid biopsies in genomics-driven oncology. Nat. Rev. Genet. 20, 71–88 (2019).

7. Forshew, T. et al. Noninvasive Identification and Monitoring of Cancer Mutations by Targeted Deep Sequencing of Plasma DNA. Sci. Transl. Med. 4, (2012).

8. Keppens, C. et al. Detection of EGFR Variants in Plasma. J. Mol. Diagn. 20, 483–494 (2018).

9. Perkins, G., Lu, H., Garlan, F. & Taly, V. Droplet-Based Digital PCR. in Advances in Clinical Chemistry vol. 79 43–91 (Elsevier, 2017).

10. Cheng, D. T. et al. Memorial Sloan Kettering-Integrated Mutation Profiling of Actionable Cancer Targets (MSK-IMPACT). J. Mol. Diagn. 17, 251–264 (2015).

11. Song, P. et al. Limitations and opportunities of technologies for the analysis of cell-free DNA in cancer diagnostics. Nat. Biomed. Eng. 6, 232–245 (2022).

12. Pantel, K. & Alix-Panabières, C. Liquid biopsy and minimal residual disease — latest advances and implications for cure. Nat. Rev. Clin. Oncol. 16, 409–424 (2019).

13. Brito-Rocha, T., Constâncio, V., Henrique, R. & Jerónimo, C. Shifting the Cancer Screening Paradigm: The Rising Potential of Blood-Based Multi-Cancer Early Detection Tests. Cells 12, 935 (2023).

14. Nirmaladevi, R. Epigenetic alterations in cancer. Front. Biosci. 25, 1058–1109 (2020).

15. Kanwal, R. & Gupta, S. Epigenetic modifications in cancer. Clin. Genet. 81, 303–311 (2012).

16. Cain, J. A., Montibus, B. & Oakey, R. J. Intragenic CpG Islands and Their Impact on Gene Regulation. Front. Cell Dev. Biol. 10, 832348 (2022).

17. Kulis, M., Queirós, A. C., Beekman, R. & Martín-Subero, J. I. Intragenic DNA methylation in transcriptional regulation, normal differentiation and cancer. Biochim. Biophys. Acta BBA - Gene Regul. Mech. 1829, 1161–1174 (2013).

18. Ibrahim, J., Peeters, M., Van Camp, G. & Op De Beeck, K. Methylation biomarkers for early cancer detection and diagnosis: Current and future perspectives. Eur. J. Cancer 178, 91–113 (2023).

19. Klein, E. A. et al. Clinical validation of a targeted methylation-based multi-cancer early detection test using an independent validation set. Ann. Oncol. 32, 1167–1177 (2021).

20. Kurdyukov, S. & Bullock, M. DNA Methylation Analysis: Choosing the Right Method. Biology 5, 3 (2016).

21. Encyclopedia of Cancer. (Elsevier, Academic Press, Amsterdam Boston Heidelberg London, 2019).

22. Werner, B. et al. Circulating cell-free DNA from plasma undergoes less fragmentation during bisulfite treatment than genomic DNA due to low molecular weight. PLOS ONE 14, e0224338 (2019).

23. Sina, A. A. I. et al. Epigenetically reprogrammed methylation landscape drives the DNA self-assembly and serves as a universal cancer biomarker. Nat. Commun. 9, 4915 (2018).

24. Anand, S., Swami, P., Goel, G. & Gupta, S. Zwitterions for impedance spectroscopy: The new buffers in town. Anal. Chim. Acta 1166, 338547 (2021).

25. Genovese, G. et al. Clonal Hematopoiesis and Blood-Cancer Risk Inferred from Blood DNA Sequence. N. Engl. J. Med. 371, 2477–2487 (2014).

26. Hinrichsen, T., Dworniczak, J. K., Wachter, O., Dworniczak, B. & Dockhorn-Dworniczak, B. Detection and characterization of circulating cell free tumor DNA in cancer patients with malignant solid tumors. Liquid biopsy: a new tool in molecular pathology? LaboratoriumsMedizin 40, 313–322 (2016).

27. Gupta, S., Kilpatrick, P. K., Melvin, E. & Velev, O. D. On-chip latex agglutination immunoassay readout by electrochemical impedance spectroscopy. Lab. Chip 12, 4279 (2012).

28. Kratochvílová, I. et al. Conductivity of natural and modified DNA measured by scanning tunneling microscopy. The effect of sequence, charge and stacking. Biophys. Chem. 138, 3–10 (2008).

29. Peled, M. et al. Cell-free DNA concentration in patients with clinical or mammographic suspicion of breast cancer. Sci. Rep. 10, 14601 (2020).

30. Boscolo-Rizzo, P. et al. TERT promoter mutations in head and neck squamous cell carcinoma: A systematic review and meta-analysis on prevalence and prognostic significance. Oral Oncol. 140, 106398 (2023).

31. Hisano, O., Ito, T. & Miura, F. Short single-stranded DNAs with putative non-canonical structures comprise a new class of plasma cell-free DNA. BMC Biol. 19, 225 (2021).

32. Daraghma, S. M. A., Talebi, S. & Periasamy, V. Understanding the electronic properties of single- and double-stranded DNA. Eur. Phys. J. E 43, 40 (2020).

33. Liu, J. et al. Genome-wide cell-free DNA methylation analyses improve accuracy of non-invasive diagnostic imaging for early-stage breast cancer. Mol. Cancer 20, 36 (2021).

34. Erichsen, L., Thimm, C. & Santourlidis, S. Methyl Group Metabolism in Differentiation, Aging, and Cancer. Int. J. Mol. Sci. 23, 8378 (2022).

35. Lakshminarasimhan, R. & Liang, G. The Role of DNA Methylation in Cancer. in DNA Methyltransferases - Role and Function (eds. Jeltsch, A. & Jurkowska, R. Z.) vol. 945 151–172 (Springer International Publishing, Cham, 2016).

36. Khandelwal, A. V. et al. AC Conductivity Measurements of Ultradilute Colloidal Suspensions in HEPES Buffer. Langmuir 35, 14725–14733 (2019).

37. Lapin, M. et al. Fragment size and level of cell-free DNA provide prognostic information in patients with advanced pancreatic cancer. J. Transl. Med. 16, 300 (2018).

38. Genereux, J. C. & Barton, J. K. Mechanisms for DNA Charge Transport. Chem. Rev. 110, 1642–1662 (2010).

39. Prasher, R., Phelan, P. E. & Bhattacharya, P. Effect of Aggregation Kinetics on the Thermal Conductivity of Nanoscale Colloidal Solutions (Nanofluid). Nano Lett. 6, 1529–1534 (2006).

40. Wilson, M. S. Electrochemical Immunosensors for the Simultaneous Detection of Two Tumor Markers. Anal. Chem. 77, 1496–1502 (2005).

41. Ma, H. et al. An impedance-based integrated biosensor for suspended DNA characterization. Sci. Rep. 3, 2730 (2013).

42. McDonald, J. C. et al. Fabrication of microfluidic systems in poly(dimethylsiloxane). Electrophoresis 21, 27–40 (2000).

