## Supplementary Information for "Impedance-based assay for pan-cancer early and rapid detection of cell-free DNA"

**Table S1.** Comparison of our system with various other electrical/electrochemical impedance methods reported in the literature.

| Detection method | Target | Receptor | Amplification method | Linear range | LOD | Time | Refs |
|---|---|---|---|---|---|---|---|
| Electrochemical (IS) | ctDNA | Probe DNA | None | 0.1 to 100 fM | 1 aM | - | 1 |
| Electrochemical (IS) | DNA PCR products | Receptor-less | NP labelling | - | 25–30 DNA copies | - | 2 |
| Electrochemical (DPV) | cfDNA mutation | PNA capture probe | None | 10.27 fM-1.03 nM | 10.27 fM | 15 min | 3 |
| Electrochemical (DPV) | gDNA methylation | Direct gDNA adsorption | None | - | - | 10 min | 4 |
| Electrochemical (DPV) | cfDNA | Triple-helix molecular switch | Target recycling + branched TdT | 10 aM - 1 pM | 2.4 aM | | 5 |
| Electrochemical (SWV) | ctDNA (101 nucleotides, NSCLC) | DNA-Au@MNPs | MB tagged on electrode & Au@MNPs | 200 aM - 20 nM | 5 fM | 20 min | 6 |
| Electrochemical (SWV) | ctDNA mutation & methylation | PNA & anti-5-mC Ab | AuNPs & lead phosphate apoferritin | 50-10000 fM | 10 fM | 30 min | 7 |
| Electrochemical (chronoamperometry) | Methylated oligos | Tetrahedral DNA nanostructure probe | AuNP deposition, HCR & HRP | 1 aM - 1 pM | 1 aM | 6 h | 8 |
| Electrical (IS) | DNA PCR products | None | None | - | 148 bp | - | 9 |
| Electrical (IS) [Present Work] | cfDNA methylation | None | None | - | 5.14 pM | < 5 min | - |

IS: Impedance Spectroscopy; DPV: Differential pulse voltammetry; SWV: Square-wave voltammetry; PNA; Peptide nucleic acids; gDNA: Genomic DNA; AuNP: gold nanoparticle; anti-5-mC Ab: Monoclonal anti-5-methylcytosine antibody; PCR: Polymerase chain reaction; ctDNA: circulating tumor DNA; DNA-Au@MNPs: DNA modified gold-coated magnetic NPs; HCR: hybridization chain reaction; HRP: Horseradish peroxidase; TdT: terminal deoxynucleotidyl transferase; MB: methylene blue; NSCLS; Non-small cell lung cancer

**Table S2.** Clinical information of cancer samples used for cfDNA extraction. Information where not available is left blank.

| Sample ID | Cancer | Stage | | Age (yrs) | Sex | Estimated cfDNA concentration in plasma (ng/ml) | ΔZ (Ω) | +/-2SD (Ω) |
|---|---|---|---|---|---|---|---|---|
| C1 | Brain | - | IV | 49 | M | 20.88 | -13.52 | 1.5 |
| C2 | Brain | - | IV | 47 | M | 128.88 | -11.55 | 1.9 |
| C3 | Brain | - | IV | 33 | M | 231.00 | -15.365 | 1.74 |
| C4 | Brain | - | IV | 61 | M | 255.00 | -12.82 | 0.16 |
| C5 | Breast | pT1aN0 | IA | 57 | F | 12.00 | -6 | 0.86 |
| C6 | Breast | pT1N0 | IA | 66 | F | 111.00 | -7.65 | 0.38 |
| C7 | Breast | mpT1bN0 | IA | 65 | F | 129.00 | -23.67 | 1.2 |
| C8 | Breast | pT1cN0 | IA | 66 | F | 327.00 | -23.85 | 2.54 |
| C9 | Breast | pT2N0 | IIA | 72 | F | 72.00 | -12.05 | 1.04 |
| C10 | Breast | pT2N0 | IIA | 49 | F | 93.00 | -12.86 | 0.3 |
| C11 | Breast | pT2N0M0 | IIA | 63 | F | 123.00 | -31.935 | 1.04 |
| C12 | Breast | pT2N0 | IIA | 76 | F | 156.00 | -19.95 | 0.38 |
| C13 | Breast | pT2N0 | IIA | 52 | F | 282.00 | -22.72 | 2.4 |
| C14 | Breast | ypTisN1a | IIA | 53 | F | 406.15 | -23.34 | 5.56 |
| C15 | Breast | pT2N0 | IIA | 28 | F | 726.00 | -23.57 | 0.14 |
| C16 | Breast | pT2N1 | IIB | 43 | F | 96.00 | -1.72 | 1.9 |
| C17 | Breast | cT2N1M0 | IIB | 48 | F | 152.88 | -7.52 | 0.4 |
| C18 | Breast | cT2N1M0 | IIB | 63 | F | 157.98 | -11.915 | 0.42 |
| C19 | Breast | cT2N1M0 | IIB | 55 | F | 158.28 | -23.31 | 0.02 |
| C20 | Breast | cT2N1M0 | IIB | 46 | F | 288.00 | -1.875 | 0.26 |
| C21 | Breast | pT2N1 | IIB | 68 | M | 762.00 | -31.4 | 1.24 |
| C22 | Breast | pT2N2a | IIIA | 41 | F | 42.00 | -8.52 | 1.3 |
| C23 | Breast | pT2N2a | IIIA | 31 | F | 90.00 | -18.26 | 1.76 |
| C24 | Breast | pT2N2a | IIIA | 64 | F | 342.00 | -25.22 | 0.32 |
| C25 | Breast | pT2N3a | IIIC | 55 | F | 234.00 | -21.9 | 1.04 |
| C26 | Breast | pT4bN3a | IIIC | 79 | F | 543.00 | -53.82 | 2.78 |
| C27 | Breast | pT3N3a | IIIC | 60 | F | 558.00 | -33.45 | 1.2 |
| C28 | Breast | - | - | 36 | F | 39.88 | -21.26 | 3.06 |
| C29 | Breast | - | - | 51 | F | 48.60 | -25.64 | 5.26 |
| C30 | Breast | - | - | 59 | F | 71.74 | -28.62 | 3.3 |
| C31 | Cervical | - | - | 50 | M | 1077.41 | -0.93 | 3.32 |
| C32 | Colorectal | cT3N0M0 | IIA | 76 | M | 80.70 | -9.2 | 2.54 |
| C33 | Colorectal | cT3N0M0 | IIA | 72 | M | 171.00 | -21.545 | 1.28 |
| C34 | Colorectal | cT4N1M0 | III | 53 | M | 54.00 | -0.015 | 5.76 |
| C35 | Colorectal | cT4N2M1 | IV | 57 | F | 215.40 | -15.83 | 0.42 |
| C36 | Gallbladder | GRADE II | II | 61 | M | 168.00 | -8.67 | 2.34 |
| C37 | Gallbladder | PT3N0M0 | IIIA | 60 | M | 379.56 | -13.595 | 1.6 |
| C38 | Gallbladder | PT3N1M0 | IIIB | 58 | F | 115.76 | -11.18 | 2.78 |
| C39 | Gallbladder | - | M1 | 54 | M | 105.00 | -2.46 | 1.42 |
| C40 | Gallbladder | - | M1 | 64 | F | 165.00 | -27.98 | 2.18 |
| C41 | Head & Neck | pT1N0 | I | 53 | M | 315.78 | -30.985 | 0.72 |
| C42 | Head & Neck | pT2N0 | II | 82 | F | 144.00 | -6.8 | 0.42 |
| C43 | Head & Neck | pT2pN0 | II | 46 | M | 179.88 | -13.985 | 0.9 |
| C44 | Head & Neck | T2N0M0 | II | 66 | M | 219.00 | -43.33 | 1.56 |
| C45 | Head & Neck | pT2N0 | II | 50 | F | 321.00 | -4.95 | 2.78 |
| C46 | Head & Neck | T2N0M0 | II | 50 | M | 402.00 | -12.73 | 0.08 |
| C47 | Head & Neck | pT3N0 | III | 46 | M | 150.00 | -30.245 | 2.02 |

| | | | | | | | | |
|---|---|---|---|---|---|---|---|---|
| C48 | Head & Neck | pT1N1 | III | 43 | M | 216.00 | -16.12 | 0.98 |
| C49 | Head & Neck | T4aN2aM0 | IVA | 42 | M | 78.00 | -32.365 | 1.76 |
| C50 | Head & Neck | pT4apN0 | IVA | 46 | M | 81.00 | -30.65 | 1.56 |
| C51 | Head & Neck | pT2pN2a | IVA | 59 | F | 86.40 | -33.175 | 3.16 |
| C52 | Head & Neck | T4N2cM0 | IVA | 57 | M | 120.00 | -23.36 | 1.86 |
| C53 | Head & Neck | T4aN0M0 | IVA | 46 | M | 120.00 | -33.415 | 0.1 |
| C54 | Head & Neck | pT3pN2a | IVA | 55 | F | 162.00 | -16.465 | 1.38 |
| C55 | Head & Neck | pT4aN2a | IVA | 60 | M | 318.00 | -13.01 | 1.16 |
| C56 | Head & Neck | pT4apN3b | IVB | 54 | M | 214.80 | -18.37 | 1.14 |
| C57 | Head & Neck | pT2N3b | IVB | 39 | M | 237.00 | -9.29 | 1.4 |
| C58 | Head & Neck | - | - | 33 | M | 46.38 | -26.85 | 1.84 |
| C59 | Kidney | pT1N0 | I | 56 | M | 123.00 | -3.06 | 2.72 |
| C60 | Kidney | pT1a | I | 48 | M | 230.88 | -51.595 | 2.72 |
| C61 | Kidney | pT1bN0 | I | 51 | M | 321.00 | -25.73 | 1.34 |
| C62 | Kidney | pT1bNx | I | 71 | F | 366.00 | -6.21 | 1.82 |
| C63 | Kidney | pT1Nx | I | 78 | M | 642.00 | -56.145 | 0.6 |
| C64 | Kidney | pT2bN0 | II | 56 | M | 210.00 | -6.42 | 2.28 |
| C65 | Kidney | pT2aNx | II | 65 | M | 210.00 | -29.405 | 1.72 |
| C66 | Kidney | pT2aNx | II | 69 | M | 282.00 | -22.96 | 2.3 |
| C67 | Kidney | pT3aN0 | III | 38 | M | 60.00 | -15.765 | 0.66 |
| C68 | Kidney | pT3a(ATLE AST)Nx | III | 63 | M | 111.00 | -3.23 | 1.48 |
| C69 | Kidney | pT2aN0 | III | 49 | M | 117.00 | -4.67 | 0.32 |
| C70 | Kidney | pT3aNx | III | 53 | F | 126.00 | -2.32 | 3.68 |
| C71 | Kidney | pT2aN1 | III | 45 | M | 168.00 | -5.8 | 0.02 |
| C72 | Kidney | pT2N1 | III | 72 | M | 192.00 | -26.955 | 1.38 |
| C73 | Kidney | pT3aN1 | III | 61 | M | 234.00 | -18.68 | 0.72 |
| C74 | Kidney | pT3aN0 | III | 45 | F | 255.00 | -49.165 | 1.76 |
| C75 | Kidney | pT1N1 | III | 55 | F | 276.00 | -8.15 | 1.08 |
| C76 | Kidney | pT3aN0 | III | 55 | F | 309.00 | -20.74 | 3.38 |
| C77 | Kidney | PT3N0 | III | 53 | M | 408.00 | -32.77 | 3.74 |
| C78 | Kidney | PT3Nx | III | 51 | M | 585.00 | -7.24 | 0.08 |
| C79 | Kidney | pT4N0 | IV | 36 | M | 99.00 | -31.045 | 2.42 |
| C80 | Kidney | pT4N2 | IV | 42 | M | 138.00 | -12.54 | 1.48 |
| C81 | Kidney | pT3aN1M1 | IV | 44 | M | 156.00 | -4.31 | 0.6 |
| C82 | Kidney | pT4N0 | IV | 60 | M | 171.00 | -0.4 | 0.44 |
| C83 | Kidney | pT4N0 | IV | 75 | M | 228.00 | -1.7 | 0.74 |
| C84 | Kidney | pT4N1 | IV | 64 | M | 261.00 | -23.505 | 2 |
| C85 | Kidney | pT2aNxM1 | IV | 85 | M | 366.00 | -24.95 | 1.08 |
| C86 | Kidney | pT1aNx | - | 67 | M | 210.00 | -13.45 | 1.48 |
| C87 | Kidney | - | - | - | - | 231.00 | -23.05 | 0.32 |
| C88 | Kidney | pT1aNx | - | 62 | F | 234.00 | -2.78 | 0.54 |
| C89 | Kidney | pT1aNx | - | 65 | M | 330.00 | -22.9 | 2.4 |
| C90 | Lymph | - | - | 55 | M | 35.20 | -15.01 | 2.78 |
| C91 | Lymph | - | - | 65 | M | 60.35 | -5.12 | 0.06 |
| C92 | Lymph | - | - | 60 | F | 60.80 | -13.09 | 5.34 |
| C93 | Lymph | - | - | 45 | F | 69.90 | -8.9 | 1.42 |
| C94 | Lung | - | - | - | - | 56.98 | -2 | 2.82 |
| C95 | Lung | - | - | - | - | 68.08 | -9.235 | 4.9 |
| C96 | Lung | - | - | - | - | 74.74 | -12.8 | 7.9 |
| C97 | Lung | - | - | - | - | 79.92 | -1.75 | 2.12 |
| C98 | Lung | - | - | - | - | 79.92 | -4 | 0 |
| C99 | Lung | - | - | - | - | 82.88 | -7.15 | 8.06 |
| C100 | Lung | - | - | - | - | 84.36 | -6 | 2.82 |

| | | | | | | | | |
|---|---|---|---|---|---|---|---|---|
| C101 | Lung | - | - | - | - | 89.54 | -3.5 | 1.4 |
| C102 | Lung | - | - | - | - | 95.46 | -2 | 2.82 |
| C103 | Lung | - | - | - | - | 99.90 | -3.75 | 2.12 |
| C104 | Lung | - | - | - | - | 105.8 | -31.5 | 1.4 |
| C105 | Lung | - | - | - | - | 116.18 | -3 | 0 |
| C106 | Lung | - | - | - | - | 118.40 | -3 | 0 |
| C107 | Lung | - | - | - | - | 128.76 | -29.5 | 1.4 |
| C108 | Lung | - | - | - | - | 137.64 | -1.66 | 4.12 |
| C109 | Lung | - | - | - | - | 142.08 | -3 | 5.64 |
| C110 | Lung | - | - | - | - | 145.78 | -7.85 | 3.24 |
| C111 | Lung | - | - | - | - | 155.40 | -6.35 | 0.98 |
| C112 | Lung | - | - | - | - | 167.98 | -8.6 | 1.12 |
| C113 | Lung | - | - | - | - | 174.64 | -4.4 | 4.52 |
| C114 | Lung | - | - | - | - | 176.86 | -10.7 | 3.66 |
| C115 | Lung | - | - | - | - | 184.26 | -6.25 | 7.76 |
| C116 | Lung | - | - | - | - | 202.76 | -7.8 | 5.08 |
| C117 | Lung | - | - | - | - | 223.23 | -25.2 | 9.04 |
| C118 | Ovary | - | I | 55 | F | 243.00 | -24.17 | 1.36 |
| C119 | Ovary | - | III | 63 | F | 111.00 | -9.96 | 1.74 |
| C120 | Ovary | - | IIIC | 48 | F | 183.00 | -9.49 | 2.12 |
| C121 | Ovary | - | IV | 58 | F | 174.00 | -15.81 | 2.38 |
| C122 | Ovary | - | IV | 52 | F | 249.00 | -16.5 | 2.00 |
| C123 | Pancreas | pT2N0 | IB | 65 | F | 192.00 | -17.33 | 0.9 |
| C124 | Pancreas | pT4N0 | III | 38 | M | 162.00 | -14.14 | 0.16 |
| C125 | Penis | - | - | 45 | M | 40.71 | -16.19 | 0.04 |
| C126 | Prostate | pT2N0 | I | 71 | M | 144.00 | -7.86 | 2.06 |
| C127 | Prostate | pT2N0 | I | 60 | M | 201.00 | -12.71 | 0.84 |
| C128 | Prostate | pT3bN0 | III | 69 | M | 318.00 | -22.4 | 1.62 |
| C129 | Prostate | pT3bN0 | III | 69 | M | 411.00 | -8.19 | 0.94 |
| C130 | Prostate | pT3bN1 | IV | 76 | M | 165.00 | -9.29 | 2.86 |
| C131 | Prostate | pT3bN1 | IV | 65 | M | 264.00 | -11.54 | 1.56 |
| C132 | Prostate | pT3bN1 | IV | 76 | M | 543.00 | -34.9 | 0.08 |
| C133 | Salivary gland | - | - | 36 | M | 52.41 | -8.66 | 2.1 |
| C134 | Thyroid | pT2Nx | I | 30 | F | 95.58 | -6.935 | 0.64 |
| C135 | Thyroid | pT1aNx | I | 24 | F | 238.20 | -19.735 | 1 |
| C136 | Thyroid | T3aN1b | I | 36 | F | 309.00 | -2.105 | 0.42 |
| C137 | Thyroid | T3aN1b | I | 29 | F | 390.00 | -2.84 | 1.78 |
| C138 | Thyroid | pT2N0 | II | 61 | F | 90.00 | -2.4 | 2.14 |
| C139 | Thyroid | pT3aNx | II | 63 | M | 114.00 | -6.09 | 2.26 |
| C140 | Thyroid | pT2N0 | II | 52 | F | 117.00 | -24.85 | 0.68 |
| C141 | Thyroid | pT2 | II | 29 | M | 129.00 | -3.42 | 1.36 |
| C142 | Thyroid | pT2N0 | II | 33 | M | 261.00 | -18.14 | 1.32 |

| | | | | | | | | |
|---|---|---|---|---|---|---|---|---|
| C143 | Thyroid | pT2N0 | II | 43 | F | 294.00 | -22.13 | 2.5 |
| C144 | Thyroid | pT3aN1b | III | 64 | F | 54.00 | -0.78 | 1.52 |
| C145 | Thyroid | pT3N0 | III | 54 | F | 228.00 | -1.49 | 0.12 |
| C146 | Thyroid | pT3N0 | III | 50 | F | 252.00 | -15.26 | 0.82 |
| C147 | Thyroid | pT1bN1b | IVA | 61 | F | 84.00 | -7.37 | 1.06 |
| C148 | Thyroid | pT3aN1b | IVA | 67 | M | 102.00 | -0.25 | 0.84 |
| C149 | Thyroid | pT2N1b | IVA | 26 | F | 117.00 | -14.63 | 2.1 |
| C150 | Thyroid | T3bN1b | IVA | 78 | F | 138.00 | -6.02 | 1.08 |
| C151 | Thyroid | pT3aN1a | IVA | 66 | F | 159.00 | -3.31 | 0.74 |
| C152 | Thyroid | T1aN1b | IVA | 50 | F | 213.00 | -9.67 | 1.74 |
| C153 | Unknown | - | Metastatic | 60 | M | 53.96 | -13.89 | 6.64 |
| C154 | Unknown | - | Metastatic | 45 | M | 86.93 | -23.48 | 1.52 |
| C155 | Unknown | - | Metastatic | 45 | M | 133.25 | -11.26 | 2.68 |
| C156 | Unknown | - | - | - | - | 43.29 | -1.96 | 2.68 |
| C157 | Unknown | - | - | 41 | M | 47.60 | -4.73 | 0.64 |
| C158 | Unknown | - | - | 62 | M | 51.39 | -3.3 | 8.2 |
| C159 | Unknown | - | - | 56 | M | 56.03 | -24.25 | 2.96 |
| C160 | Unknown | - | - | - | M | 56.58 | -17.95 | 3.26 |
| C161 | Unknown | - | - | 26 | F | 62.49 | -2.5 | 4.8 |
| C162 | Unknown | - | - | 50 | M | 71.50 | -24.6 | 2.84 |
| C163 | Unknown | - | - | 56 | M | 93.85 | -11.77 | 0.66 |
| C164 | Unknown | - | - | - | - | 99.90 | -3.75 | 2.12 |
| C165 | Unknown | - | - | - | - | 150.00 | -15.27 | 2.1 |
| C166 | Unknown | - | - | - | - | 396.00 | -32.06 | 0.9 |

**Table S3.** Information of healthy samples used for cfDNA extraction. Information where not available is left blank.

| **Sample ID** | **Age (yrs)** | **Sex** | **Estimated cfDNA concentration in plasma (ng/ml)** | **ΔZ (Ω)** | **+/-2SD (Ω)** |
|---|---|---|---|---|---|
| H1 | 23 | F | 42.60 | -65.83 | 0.36 |
| H2 | 23 | M | 116.33 | -45.85 | 1.70 |
| H3 | 23 | M | 93.75 | -55.1 | 4.56 |
| H4 | 23 | M | 105.00 | -36.37 | 2.70 |
| H5 | 24 | M | 22.50 | -49.38 | 0.22 |
| H6 | 24 | F | 58.50 | -67.76 | 0.74 |
| H7 | 25 | M | 74.00 | -63.05 | 3.90 |
| H8 | 25 | F | 36.10 | -52.25 | 1.50 |
| H9 | 25 | F | 57.00 | -65.08 | 0.15 |
| H10 | 25 | M | 49.50 | -46.21 | 2.24 |

| | | | | | |
|---|---|---|---|---|---|
| H11 | 26 | F | 79.11 | -57.59 | 3.72 |
| H12 | 26 | F | 76.94 | -40.28 | 0.56 |
| H13 | 26 | F | 58.31 | -52.48 | 1.55 |
| H14 | 26 | F | 40.92 | -65.83 | 0.35 |
| H15 | 27 | F | 40.57 | -63.53 | 2.44 |
| H16 | 27 | M | 69.00 | -59.11 | 1.68 |
| H17 | 27 | F | 69.75 | -78.70 | 6.20 |
| H18 | 28 | F | 25.50 | -73.21 | 2.10 |
| H19 | 29 | F | 54.60 | -77.96 | 2.32 |
| H20 | 29 | M | 44.23 | -66.70 | 0.60 |
| H21 | 29 | M | 55.68 | -34.80 | 7.60 |
| H22 | 29 | M | 69.33 | -32.25 | 1.50 |
| H23 | 29 | M | 111.00 | -35.34 | 4.54 |
| H24 | 30 | M | 57.78 | -46.00 | 4.00 |
| H25 | 30 | F | 54.18 | -36.00 | 2.00 |
| H26 | 30 | F | 35.94 | -43.50 | 5.00 |
| H27 | 30 | M | 27.00 | -68.94 | 0.32 |
| H28 | 30 | M | 78.00 | -67.38 | 4.72 |
| H29 | 32 | M | 51.81 | -56.00 | 0.00 |
| H30 | 32 | M | 129.00 | -56.79 | 0.32 |
| H31 | 32 | F | 61.50 | -52.13 | 4.76 |
| H32 | 34 | F | 60.26 | -31.59 | 3.17 |
| H33 | 36 | F | 45.56 | -56.15 | 2.30 |
| H34 | 38 | F | 40.67 | -45.40 | 0.00 |
| H35 | 47 | M | 60.00 | -42.26 | 3.28 |
| H36 | 50 | F | 60.00 | -50.78 | 2.26 |
| H37 | 53 | M | 150.00 | -21.37 | 1.73 |
| H38 | 55 | F | 116.25 | -47.70 | 1.41 |
| H39 | 57 | M | 78.00 | -48.54 | 1.59 |
| H40 | 62 | F | 130.00 | -43.51 | 2.12 |
| H41 | 66 | F | 99.60 | -39.62 | 1.74 |
| H42 | 68 | M | 68.50 | -43.94 | 3.03 |
| H43 | 75 | F | 105.00 | -40.79 | 2.21 |
| H44 | 77 | M | 78.75 | -40.705 | 4.03 |
| H45 | - | - | 35.00 | -58.23 | 0.98 |
| H46 | - | - | 37.00 | -71.45 | 8.36 |
| H47 | - | - | 49.00 | -57.9 | 4.24 |
| H48 | - | - | 57.00 | -42.5 | 3.00 |

| | | | | | |
|---|---|---|---|---|---|
| H49 | - | - | 71.00 | -57.85 | 1.47 |
| H50 | - | - | 79.00 | -65.02 | 0.21 |

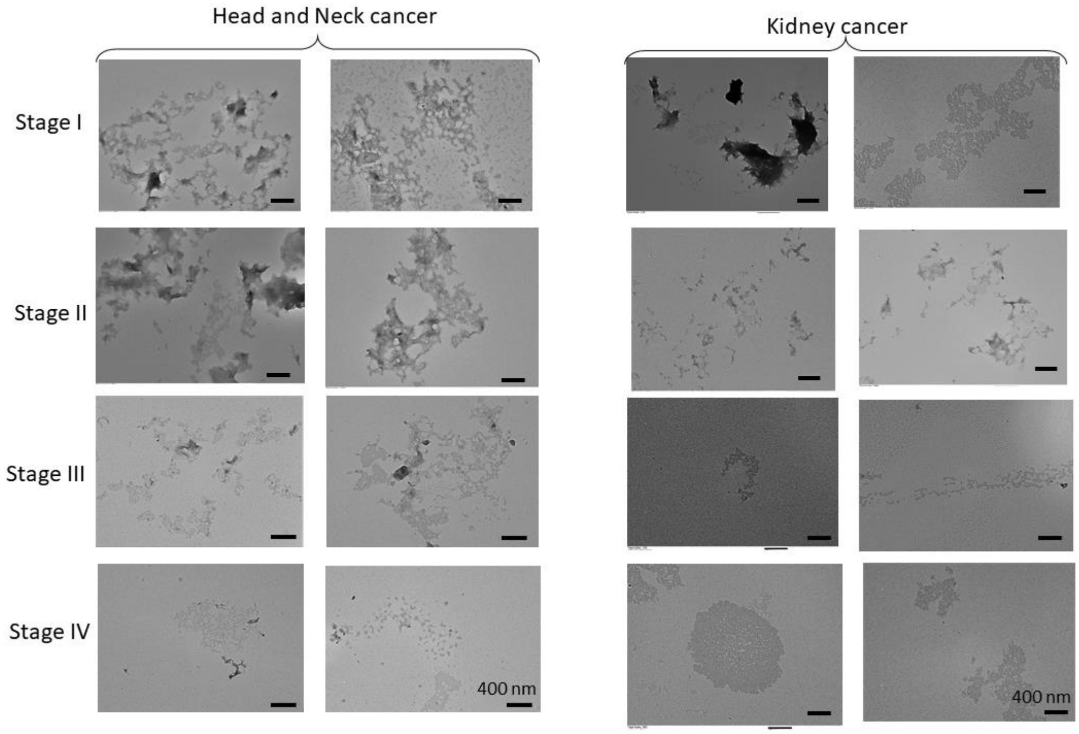


**Fig. S1.** TEM micrographs showing structural variation in cfDNA with progression of cancer stages for head and neck cancer (C41, C45, C48, C53) and kidney cancer (C62, C65, C74, C79). Sample IDs provided in the order of increasing stages. Scale bar is 400 nm.

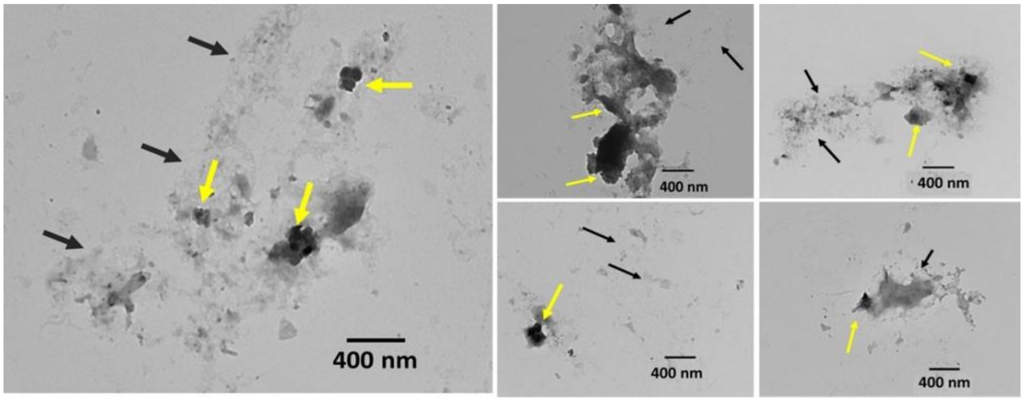


**Fig. S2** TEM micrographs of stage I breast cancer (C6) showing coexistence of aggregated (yellow arrows) and dispersed (black arrows) forms of cfDNA molecules.

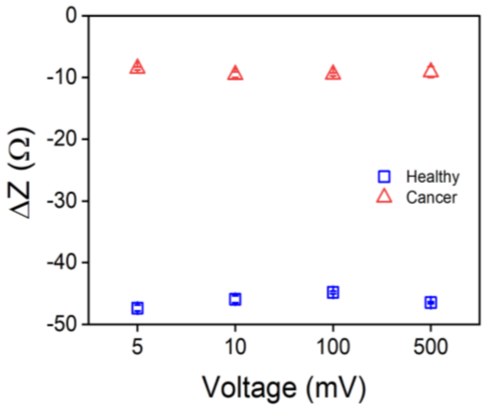


**Fig. S3.** Effect of voltage on the impedance response of healthy (H11) and cancer (C37) cfDNA samples at 100 kHz. The error bars where not visible are smaller than the data points.

**Table S4.** The effect of medium on impedance signal. The data were collected in both healthy (H25) and cancer (C104) samples. and reported as the mean of three replicates ± 2 SD.

| **Medium (pH 7.4)** | **Conductivity** (S/m) | **$\Delta Z_1$ (Ω) Healthy** | **\|S/N\|** | **$\Delta Z_2$ (Ω) Cancer** | **\|S/N\|** | **$\Delta Z = \Delta Z_2 - \Delta Z_1$ (Ω)** | **\|S/N\|** |
|---|---|---|---|---|---|---|---|
| 15 mM HEPES | 0.04 | -33.20 ± 2.80 | 11.85 | -2.00 ± 1.40 | 1.42 | -31.20 ± 3.13 | **9.96** |
| 15 mM POPSO | 0.21 | -0.59 ± 0.23 | 2.56 | 1.44 ± 0.05 | 28.8 | 2.03 ± 0.24 | **8.45** |
| 15 mM EPPS | 0.06 | -7.02 ± 1.41 | 4.97 | 9.22 ± 1.69 | 5.45 | 16.24 ± 2.20 | **7.38** |
| DI water | 4.37e-4 | (-3.05 ± 0.68)e3 | 4.48 | (2.51 ± 0.57)e3 | 4.42 | (-5.56 ± 0.88)e3 | **6.27** |
| Milli-Q water | 5.41e-4 | (-7.99 ± 1.24)e3 | 6.44 | (0.17 ± 1.46)e3 | 0.12 | (8.16 ± 1.91)e3 | **4.27** |
| 10 mM tris-EDTA | 0.11 | 1.89 ± 0.37 | 5.10 | -2.40 ± 0.94 | 2.55 | -4.29 ± 1.01 | **4.25** |
| 15 mM PIPES | 0.33 | -3.44 ± 1.04 | 3.31 | 0.04 ± 0.60 | 0.07 | -3.40 ± 1.20 | **2.83** |
| 10 mM PBS | 1.75 | 0.05 ± 0.02 | 2.50 | 0.39 ± 0.64 | 0.61 | 0.34 ± 0.64 | **0.53** |

**Sample calculation:**

Impedance of healthy sample, $Z_1$ = -33.20 ± 2.80 Ω

Impedance of cancer sample, $Z_2$ = -2.00 ± 1.40 Ω

Impedance signal change, $\Delta Z = Z_2 - Z_1$ = -33.20 – (-2.00) = -31.20 ± 3.13

(Note: Standard deviation calculation = $\sqrt{\sigma_{Z1}^2 + \sigma_{Z2}^2} = \sqrt{2.8^2 + 1.4^2}$ = 3.13)

Therefore, signal to noise ratio, S/N = 31.20/3.13 = 9.96

**Table. S5.** Effect of annealing on the overall assay response.

| S. No. | Sample id | Concentration (ng/ml) | Before annealing ($\Delta Z_1$ in Ω) | +/-2SD (Ω) | After annealing ($\Delta Z_2$ in Ω) | +/-2SD (Ω) |
|---|---|---|---|---|---|---|
| 1. | H08 | 36.1 | 61.10 | 4.10 | 109.50 | 0.71 |
| 2. | H09 | 57 | 51.50 | 2.83 | 285.50 | 0.71 |
| 3. | H15 | 40.57 | 45.40 | 6.22 | 104.65 | 0.49 |
| 4. | H19 | 54.6 | 66.40 | 0.56 | 245.50 | 0.71 |
| 5. | H29 | 51.81 | 41.0 | 1.41 | 244.95 | 7.03 |
| **Average Signal for Healthy, $\Delta Z_{Havg}$** | | | **53.08** | **3.62** | **198.02** | **3.18** |
| 6. | C96 | 74.74 | 10.00 | 3.95 | 61.00 | 1.41 |
| 7. | C97 | 79.92 | 1.75 | 1.06 | 55.85 | 1.63 |
| 8. | C98 | 79.92 | 4.00 | 0 | 41.98 | 39.19 |
| 9. | C100 | 84.36 | 6.00 | 1.41 | 14.50 | 0.71 |
| 10. | C106 | 118.4 | 3.00 | 0 | 55.05 | 16.05 |
| 11. | C115 | 184.26 | 6.25 | 3.89 | 78.00 | 8.48 |
| **Average Signal for cancer, $\Delta Z_{Cavg}$** | | | **5.16** | **2.37** | **51.06** | **17.65** |
| **Change in Signal between Healthy & Cancer, $\lvert\Delta Z_{Havg} - \Delta Z_{Cavg}\rvert$** | | | **47.92** | **4.32** | **146.96** | **17.93** |

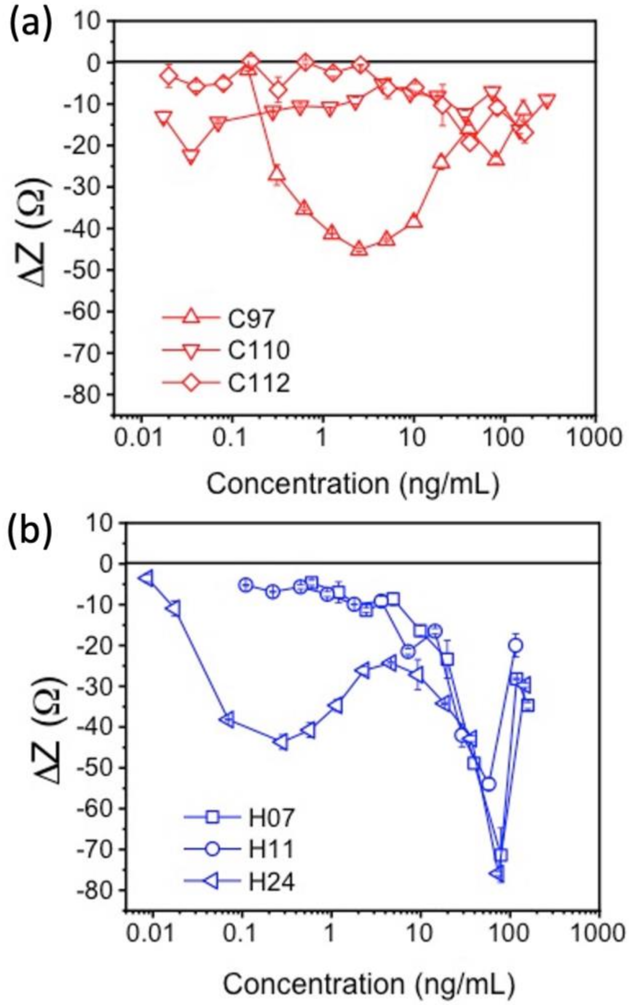


**Fig. S4.** Fig. 2E re-plotted on a logarithmic scale along the X-axis. The data in **(a)** is for cancerous cfDNA samples and in **(b)** for healthy cfDNA samples. The errors bars where not visible are smaller than the data points.

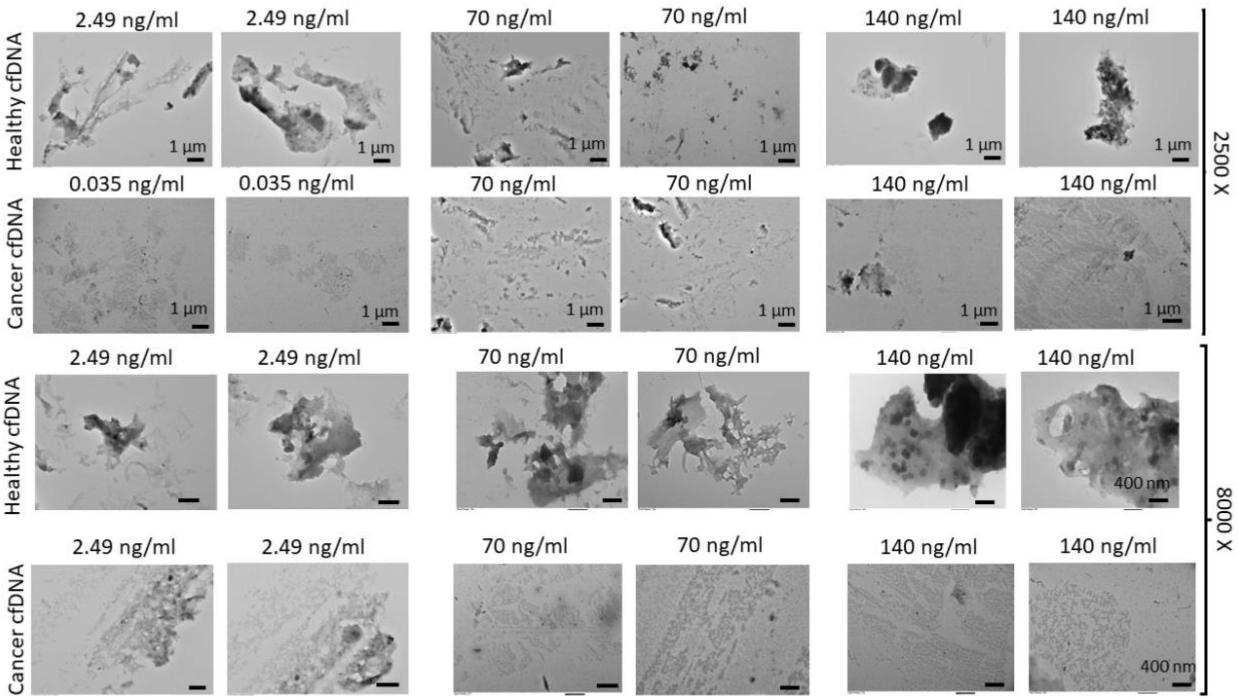


**Fig. S5** TEM micrographs of healthy (H11) and lung cancer (C98) cfDNA samples at 2500X (scale bar is 1 µm) and at 8000X (scale bar is 400 nm).

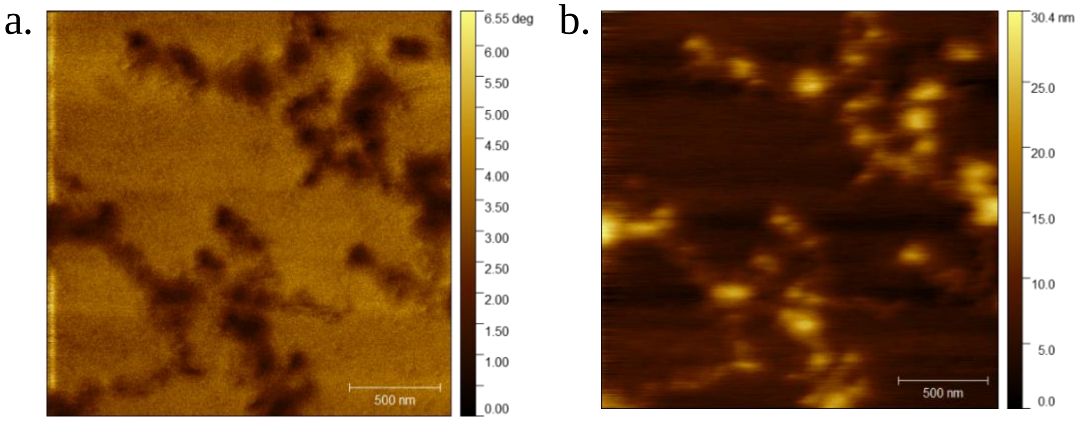


**Fig. S6 AFM characterization of healthy cfDNA.** AFM images showing the change in phase **(a)** and topography **(b)** of aggregated healthy cfDNA. The samples were prepared by drying 20 µl of cfDNA test sample on a glass coverslip and then imaging it using an ICSPI Redux AFM in ambient air tapping-mode using a diamond-like carbon (DLC) tip with less than 20 nm tip radius. The sample was scanned at a resolution of 512 pixels x 512 pixels. The scale bar indicates 500 nm.

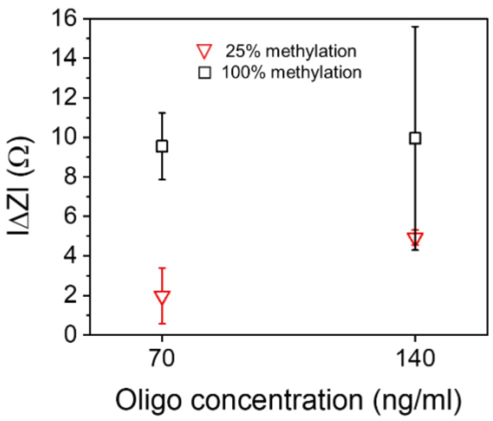


**Fig. S7** Effect of ssDNA concentration on impedance signal for 25% and 100% methylated oligos.

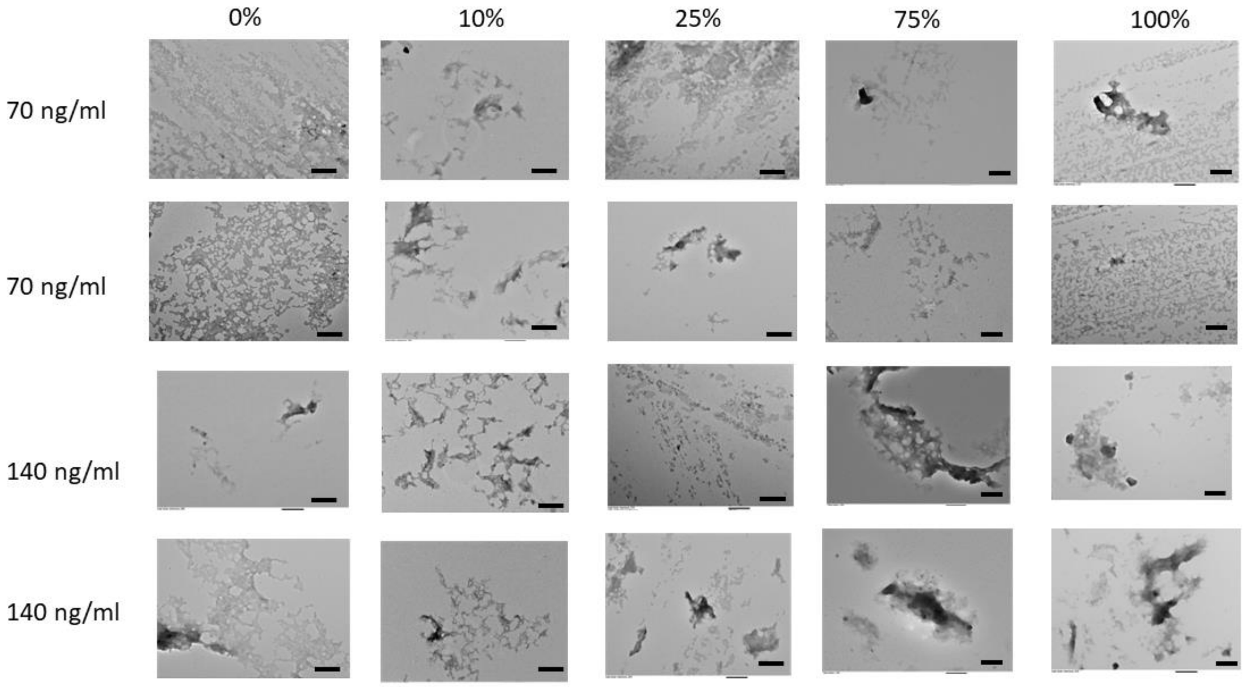


**Fig. S8** TEM micrographs of 0% to 100% methylated dsDNA at two different concentrations (scale bar is 400 nm).

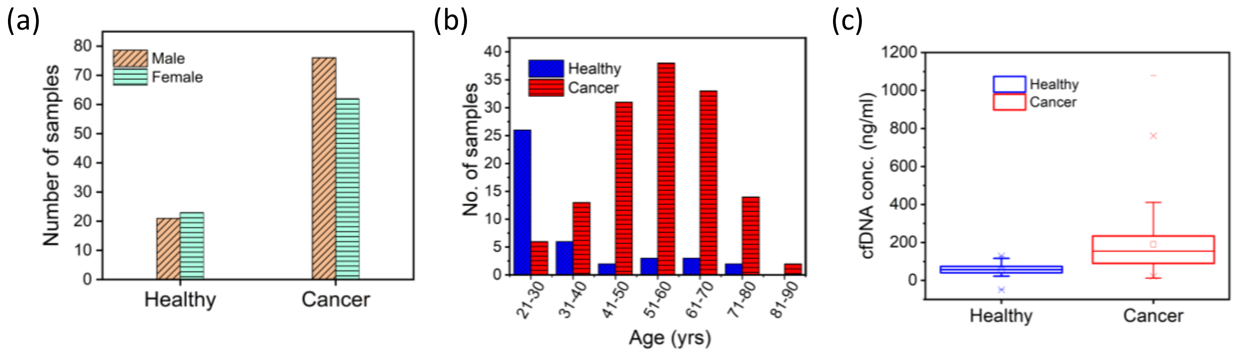


**Fig. S9.** Distribution of cancer and healthy cfDNA samples based on **(a)** sex, **(b)** age and **(c)** cfDNA concentration. In (c), the data are depicted using boxplots in which the upper border of the box indicates the upper quartile (75$^{th}$ percentile) while the lower border indicates the lower quartile (25$^{th}$ percentile), and the horizontal line in the box is the median.

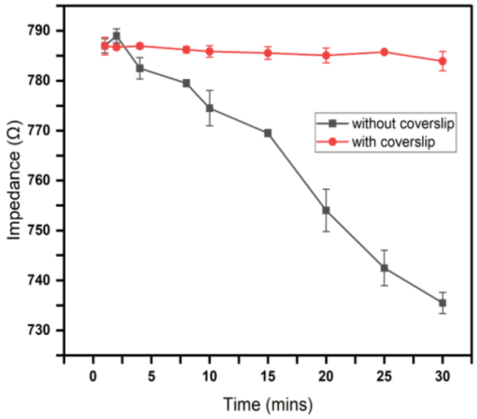


**Fig. S10.** Effect of coverslip on the time-dependent impedance response of reference solution (15 mM HEPES pH 7.4 with 2% v/v elution buffer).

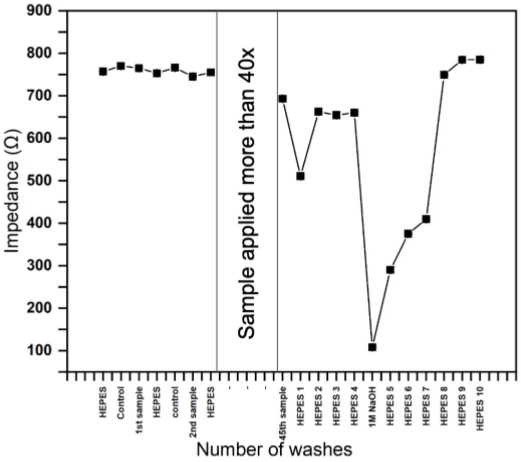


**Fig. S11.** Effect of electrode washing on electrode restoration. The electrodes could be reused up to ~ 50x with intermediate HEPES washing steps. To remove cfDNA binding beyond this point, electrodes could be completely restored after washing with 1 M NaOH.

**<u>Sample calculation for cfDNA test sample preparation</u>**

4 ml whole blood → 2 ml plasma → 20 µl elution buffer with purified cfDNA (stock)

Concentration of stock measured using NanoDrop is in ng/µl. Therefore, concentration of cfDNA estimated in plasma (assuming no loss) is,

$$\textit{cfDNA conc. in plasma (ng/ml)} = \frac{\textit{Conc. of cfDNA obtained from nanodrop (ng/}\mu l\textit{)}}{\textit{Volume of plasma (ml)}} \times 20\ \mu l$$

where, volume of plasma = 2 ml. Therefore,

$$\textit{Volume required from stock (}\mu l\textit{)} = \frac{\textit{Conc. of cfDNA calculated in plasma (ng/ml)}}{\textit{Conc. of stock solution (ng/}\mu l\textit{)}} \times 0.1\ ml$$

Here, 0.1 ml is the volume of the final test sample ($V_{test}$) prepared for measurement. It is important to note that the volume required from stock ($V_{stock}$) should always be < 2 µl. This is an important consideration to keep the background conductivity the same in all solutions. Therefore,

**$V_{test}$ |µl| = $V_{stock}$ |µl| + (2 - $V_{stock}$) |µl| elution buffer + 98 |µl| 15 mM HEPES at pH 7.4**

**References:**


1. Chu, Y., Cai, B., Ma, Y., Zhao, M., Ye, Z., & Huang, J. Highly sensitive electrochemical detection of circulating tumor DNA based on thin-layer MoS2/graphene composites. *RSC Advances 6*(27), 22673–22678 (2016).
2. Bhatt, G., Mishra, K., Ramanathan, G., & Bhattacharya, S. Dielectrophoresis assisted impedance spectroscopy for detection of gold-conjugated amplified DNA samples. *Sensors and Actuators. B, Chemical 288*, 442–453 (2019).
3. Das, J., Ivanov, I., Montermini, L. *et al.* An electrochemical clamp assay for direct, rapid analysis of circulating nucleic acids in serum. *Nature Chem* 7, 569–575 (2015).
4. Sina, A.A.I., Carrascosa, L.G., Liang, Z. *et al.* Epigenetically reprogrammed methylation landscape drives the DNA self-assembly and serves as a universal cancer biomarker. *Nat Commun* 9, 4915 (2018).
5. Wang, H., Ma, R., Sun, F., Jia, L., Zhang, W., Shang, L., Xue, Q., & Jia, W. A versatile label-free electrochemical biosensor for circulating tumor DNA based on dual enzyme assisted multiple amplification strategy. *Biosensors & Bioelectronics 122*, 224–230 (2018).
6. Chen, D., Wu, Y., Hoque, S., Tilley, R. D., & Gooding, J. J. Rapid and ultrasensitive electrochemical detection of circulating tumor DNA by hybridization on the network of gold-coated magnetic nanoparticles. *Chemical Science*, *12*(14), 5196–5201 (2021).
7. Cai, C., Guo, Z., Cao, Y., Zhang, W., & Chen, Y. A dual biomarker detection platform for quantitating circulating tumor DNA (ctDNA). *Nanotheranostics (Sydney, NSW. Online) 2*(1), 12–20 (2018).
8. Chen, X.; Huang, J.; Zhang, S.; Mo, F.; Su, S.; Li, Y.; Fang, L.; Deng, J.; Huang, H.; Luo, Z.; Zheng, J. Electrochemical Biosensor for DNA Methylation Detection through Hybridization Chain-Amplified Reaction Coupled with a Tetrahedral DNA Nanostructure. ACS Appl. Mater. Interfaces 11, 3745−3752 (2019).
9. Ma, H., Wallbank, R., Chaji, R. *et al.* An impedance-based integrated biosensor for suspended DNA characterization. *Sci Rep* 3, 2730 (2013).
